# No evidence for a common blood microbiome based on a population study of 9,770 healthy humans

**DOI:** 10.1101/2022.07.29.502098

**Authors:** Cedric C.S. Tan, Karrie K.K. Ko, Hui Chen, Jianjun Liu, Marie Loh, SG10K_Health Consortium, Minghao Chia, Niranjan Nagarajan

**Author notes:** Correspondence (C.C.S. Tan) or (N. Nagarajan). These authors jointly supervised this work (Minghao Chia, Niranjan Nagarajan).

## Abstract

Human blood is conventionally considered sterile. Recent studies have challenged this, suggesting the presence of a blood microbiome in healthy humans. We present the largest investigation to date of microbes in blood, based on shotgun sequencing libraries from 9,770 healthy subjects. Leveraging the availability of data from multiple cohorts, we stringently filtered for laboratory contaminants to identify 117 microbial species detected in the blood of sampled individuals, some of which had signatures of DNA replication. These primarily comprise of commensals associated with human body sites such as the gut (*n*=40), mouth (*n*=32), and genitourinary tract (*n*=18), which are species that are distinct from common pathogens detected in clinical blood cultures based on more than a decade of records from a tertiary hospital. Contrary to the expectations of a shared blood microbiome, no species were detected in 84% of individuals, while only a median of one microbial species per individual was detected in the remaining 16%. Futhermore, microbes of the same species were detected in <5% of individuals, no co-occurrence patterns similar to microbiomes in other body sites was observed, and no associations between host phenotypes (e.g. demographics and blood parameters) and microbial species could be established. Overall, these results do not support the hypothesis of a consistent core microbiome endogenous to human blood. Rather, our findings support the transient and sporadic translocation of commensal microbes, or their DNA, from other body sites into the bloodstream.

## Introduction

In recent years, there has been considerable interest regarding the existence of a microbiome in the blood of healthy individuals, and its links to health and disease. Human blood is traditionally considered a sterile environment (i.e., devoid of viable microbes), where the occasional entry and proliferation of pathogens in blood can trigger a dysregulated host response, resulting in severe clinical sequelae such as sepsis, septic shock or death^1^. Asymptomatic transient bacteraemia (i.e., bacterial presence in blood) in blood donors is also known to be a major cause of transfusion-related sepsis^2^. Recent studies have suggested the presence of a blood microbiome, providing evidence for microbes circulating in human blood for healthy individuals^3–7^ (reviewed in Castillo *et al*^8^). However, most of these studies were either done in relatively small cohorts or lacked rigorous checks to distinguish true biological measurements from different sources of contamination^8^. As such, the concept of a microbial community present in the blood of healthy individuals remains controversial and is an area of active research. In this work, we analysed blood DNA sequencing data from a population study of healthy individuals, comprising of multiple cohorts processed by different laboratories with varied sequencing kits. By leveraging the large dataset (*n*=9,770) complete with batch information in our systematic differential analyses for potential contaminants, our aim was to determine whether a blood microbiome truly exists in the general population.

For meaningful discourse, it is useful to formalise what the presence of a hypothetical ‘blood microbiome’ entails. Berg et al.^9^ concluded that the term microbiome should refer to a community of microbes that interact with each other and with the environment in their ecological niche, which in our context is human blood. Therefore in a blood microbiome, the presence of microbial cells in blood from healthy individuals should exhibit community structures indicated by co-occurrence or mutual exclusion of species^10^ as seen in the microbiomes of other sites such as the gut^11^ or mouth^12^. Furthermore, we may expect the presence of core microbial species, which can be defined as species that are frequently observed and shared across individuals^13,14^, such as *Staphylococcus epidermidis* on human skin^15^. More precisely, taxa that are found in a substantial fraction of samples from distinct individuals (i.e. with high prevalence) may be considered ‘core’. Notably, the prevalence threshold for defining core taxa is arbitrary, with previous microbiome studies using values ranging from 30-100% and many of these studies opting for 100%^14^. Regardless, identifying core microbes in blood would form the basis for associating microbiome changes with human health.

Existing studies have provided evidence for the presence of microbes in the blood of healthy individuals using both culture-based^3,4^ and culture-independent^5–7^ approaches. The former approach involves blood culture experiments while the latter involves one or a combination of the following molecular methods: 16S ribosomal RNA (rRNA) quantitative polymerase chain reaction (qPCR), 16S rRNA amplicon sequencing, and shotgun sequencing of RNA or DNA. Depending on the study design, these results should be interpreted with caution due to several methodological and technical limitations which include small sample sizes, limited taxonomic resolution, difficulties in distinguishing cell-free microbial DNA from live microbial cells, and the ubiquity of environmental contamination^8,16–19^. In particular, contaminating DNA must be accounted for in order to characterize the blood microbiome. The workflow of sample processing, from skin puncture during phlebotomy, to microbial detection, is rife with opportunities for microbes or microbial nucleic acids to be introduced. Contaminating microbial cells introduced due to poor aseptic technique or insufficient disinfection of the skin puncture site^20^ affects both culture-dependent and culture-independent approaches. Sequencing-based approaches are especially sensitive to contaminant microbial DNA native to laboratory reagent kits (i.e., the ‘kitome’)^19^, exacerbated by the low microbial biomasses in blood, accompanied by high host background which increases the noise-to-signal ratio^17^. Correspondingly, comprehensive profiling of the breadth and prevalence of microbial species in blood after accounting for external sources of contamination has not yet been done and several aspects of the ‘blood microbiome’ remain unclear. For instance, are the detected microbes endogenous to blood or translocated from other body sites? Is there a core set of microbes that circulates in human blood? Is there a microbial community whose structure and function could influence host health?

To address these questions, we performed the largest scale analysis of a blood sequencing dataset to date, based on DNA libraries for 9,770 healthy individuals from six distinct cohorts (**Supplementary Table 1**). We applied various bioinformatic techniques to differentiate DNA signatures of microbes in blood from potential reagent contaminants and sequence analysis artefacts, leveraging the differences in reagent kits used to process each cohort. We detected 117 microbial species in the blood of these healthy individuals, most of which are commensals associated with the microbiomes of other body sites. Additionally, we identified DNA signatures of replicating bacteria in blood using coverage-based peak-to-trough ratio analyses^21,22^, providing a culture-independent survey that has not been achieved previously. Despite this, we found no evidence for microbial co-occurrence relationships, core species, or associations with host phenotypes. These findings challenge the paradigm of a ‘blood microbiome’ and instead support a model whereby microbes from other body sites (e.g. gut, oral) sporadically translocate into the bloodstream of healthy individuals, albeit more commonly than previously assumed. Overall, our observations serve to establish a much needed baseline for the use of clinical metagenomics in investigating bloodstream infections.

## Results

### Robust inference of microbial DNA signatures in blood based on multi-cohort analysis

Blood samples from healthy individuals typically contain low microbial biomass accompanied by high host DNA background^17^, making it difficult to discriminate between biologically relevant signals from artefactual ones. We first addressed artefacts arising during bioinformatic sequence analysis by performing stringent quality control on samples (**Figure 1a**), comprising of read quality trimming and filtering, removal of low complexity sequences that are of ambiguous taxonomic origin, exclusion of reads that likely originate from human DNA (**Methods**), and removal of samples with low number of reads (<100 read pairs) of microbial origin after taxonomic classification with *Kraken2*^23^. This provided a species-level characterisation of microbial DNA signatures in blood for most (*n*=8,892) samples. To minimise noise due to false positive taxonomic assigments, we applied an abundance-cutoff based filter to discriminate between species that are likely present from those that could be misclassification artefacts (**Methods**). Additionally, we validated the reliability of the microbial species detected via *Kraken2* by comparison to read alignment analysis using reference genomes, where recovery of large fractions of a microbial reference covered uniformly by mapped reads improves our confidence that they are true positives as opposed to sequencing or analysis artefacts^24,25^. We validated 96% of the microbial species that had sufficient read coverage using this mapping-based approach. We further observed an excellent linear relationship between the number of *Kraken2*-assigned read pairs and the number of aligned read pairs on the log10 scale (slope=1.15; *F*=154, *d*.*f*.=1, *p*<0.001; **Supplementary Figure 1**), suggesting that *Kraken2* taxonomic assignments are a reliable proxy for the more precise and stringent read alignment approach. These findings collectively provide confidence that the microbial species detected in our blood sequencing libraries are not likely sequence analysis artefacts.

**Figure 1:**
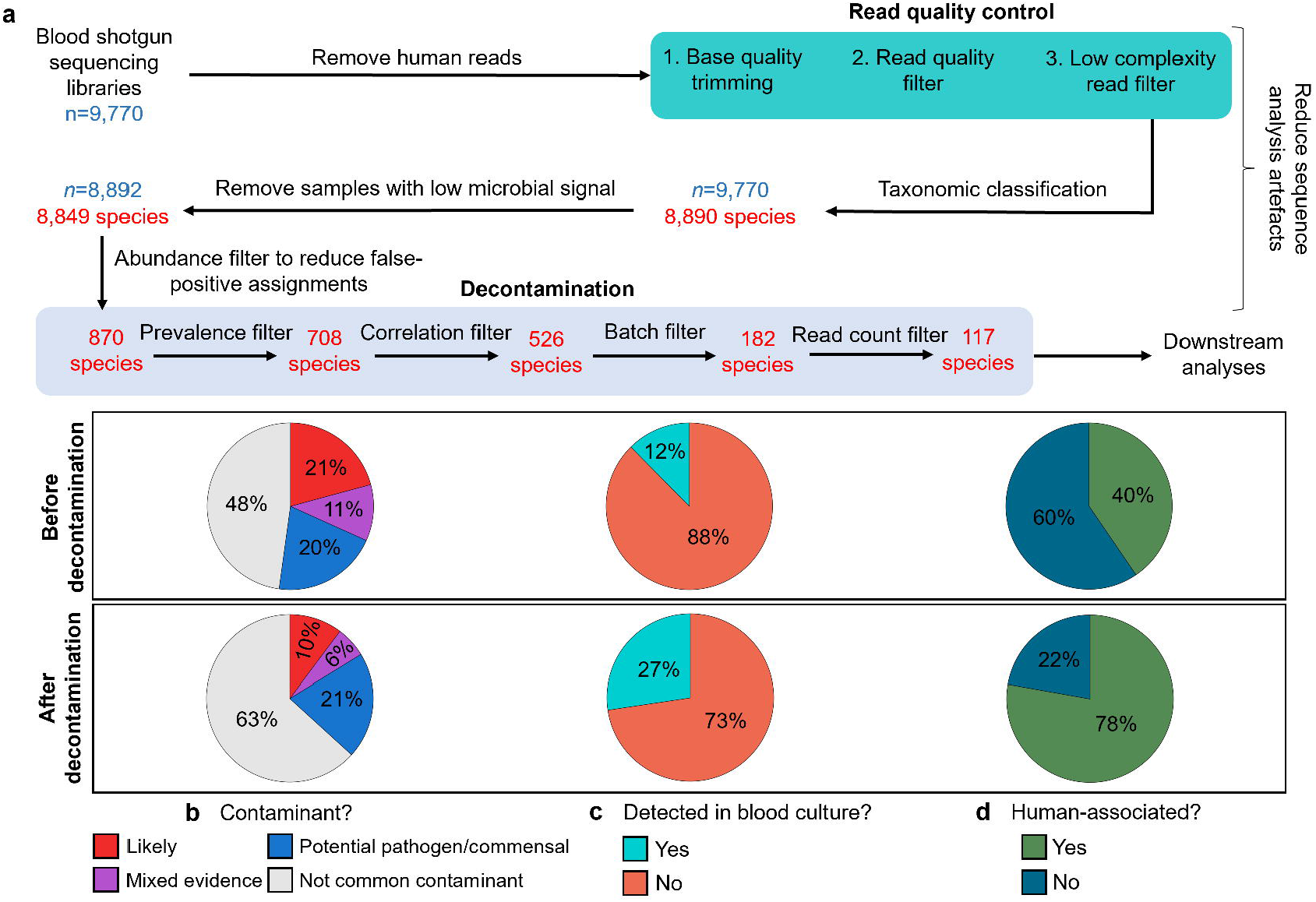
Decontamination results. (a) Summary of pre-processing steps and filters applied to taxonomic profiles (*n*=9,770 individuals) and the number of species retained after each filter. Pie charts showing the proportion of microbial species that are (b) human-associated, (c) common sequencing contaminants, and (d) detected in blood culture records, before and after applying the decontamination filters.

To address artefactual signals arising due to reagent and handling contamination during sample processing, we used a series of stringent decontamination filters (**Figure 1a**). These filters are based on the idea that contamination artefacts will lead to false positive detections that are often correlated with each other (within-batch consistency) and biased towards specific laboratory batches (between-batch variability; **Supplementary Figure 2**)^26^, and such analysis was found to be highly effective for *in silico* decontamination in previous studies^27–29^ (**Methods**). Additionally, the identification of batch-specific contaminants in this study was greatly aided by the availability of multiple large cohorts of healthy individuals (**Supplementary Table 1**), and corresponding rich batch information, including reagent kit types and lot numbers. Application of reagent and handling contamination filters resulted in a final list of 117 microbial species that were detected in the whole blood samples of 8,892 individuals (**Supplementary Table 2**). The list of 117 confidently detected microbial species spanned 56 genera, and comprised of 110 bacteria, 5 viruses and 2 fungi.

To estimate the effectiveness of our filtering strategy in improving biological signal while reducing contamination noise, we examined the types of microbial species detected in our dataset before (870 species) and after (117 species) all filters were applied (**Figure 1b-d**). Firstly, the microbial species were cross-referenced against a published list of common genera seen as contaminants in sequencing data as curated by Poore *et al*^30^ and derived from the list published by Salter *et al*^19^. In this list, genera were either classified as likely contaminants, mixed-evidence (i.e., both a pathogen and common contaminant), or potential pathogens/commensals. Following decontamination, the proportion of detected species that are classified as contaminants decreased from 21% to 10% (**Figure 1b**). Next, the microbial species were compared against human blood culture records spanning more than a decade (2011-2021) from a tertiary hospital (**Figure 1c**). These blood cultures were typically ordered if clinical indications of bacteraemia were present, and therefore represent the range of microbial species that are known to cause symptomatic infection as detected in a clinical setting. The proportion of species that have been cultured from blood increased from 12% to 27% after decontamination, suggesting that our filtering procedures enriched for microbial species which are capable of invading the bloodstream. Finally, we compared the proportion of human-associated microbes before and after decontamination using a host-pathogen association database describing the host range of pathogens^31^ (**Figure 1d**). For species that were not found in this database, a systematic PubMed search (**Methods**) was performed to determine if there was at least one past report of human infection. The proportion of human-associated species increased from 40% to 78% after decontamination, indicating that they are more likely to be biologically relevant. Finally, we tested our results against the null hypotheses that the 117 microbial species retained after decontamination produced the same proportions of species classified as likely contaminants, human-associated, or that were detected in blood culture compared to species picked at random (**Methods**). This analysis suggests that our decontamination filters significantly decreased the proportions of likely contaminants, while increasing the proportions of human-associated species and those detected in blood cultures (*p*<0.005; **Supplementary Figure 3**). These results collectively suggest that by using a set of contaminant-identification heuristics, our filters are sensitive and specific in retaining a higher proportion of biologically relevant taxa while removing likely contaminants.

### Blood microbial signatures from healthy individuals reflect sporadic translocation of DNA from commensals

We next determined the fraction of distinct, healthy individuals for which microbes could be detected (i.e., prevalence). Notably, the most prevalent microbial species, *C. acnes*, was observed in 4.7% of individuals (**Figure 2a**), suggesting that none of the 117 microbes can be considered ‘core’ species that are consistently detected across most healthy individuals. Additionally, we did not detect any microbial species in most (82%) of the samples after decontamination (**Figure 2b**), whereas the remaining 18% of samples had a median of only one microbial species per sample. This low number of species detected per sample was not due to insufficient sequencing depth since there was a weak negative correlation between the number of confidently detected species per sample and the microbial read depth (Spearman’s ρ=-0.232, *p*<0.001). Furthermore, some samples containing no microbial species had a microbial read count of up to ∼2.1 million (median=6,187 reads; distribution shown in **Supplementary Figure 4**). That is, even though a considerable number of reads were classified as microbial, they were all assigned to contaminant species. These results suggest that the presence of microbes in the blood of healthy and apparently asymptomatic individuals, as estimated by our detection methods, is infrequent and sporadic.

**Figure 2:**
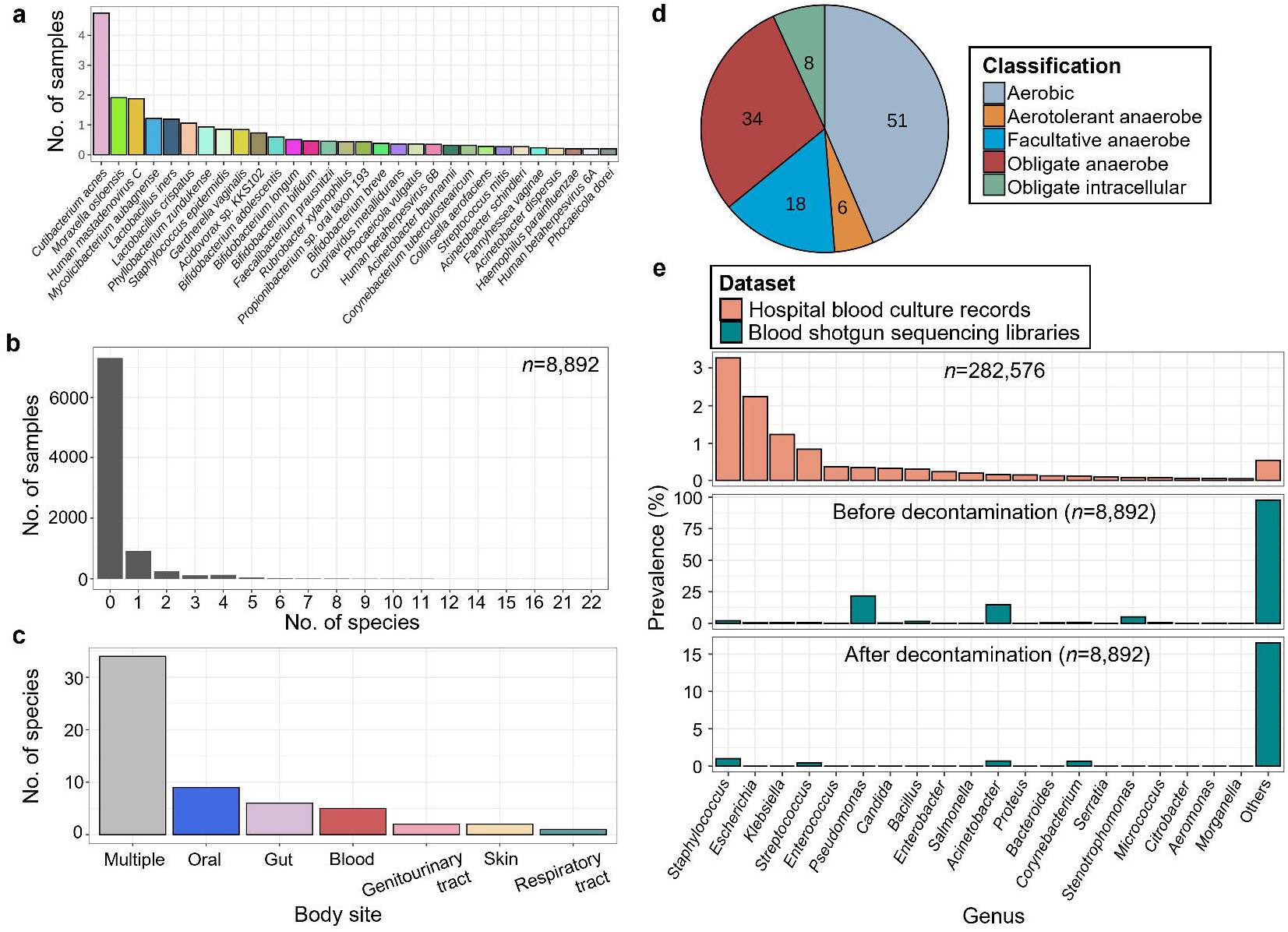
Microbial signatures in human blood from healthy individuals. (a) Bar chart showing the prevalence of the top 50 confidently detected microbial species in all 8,892 blood sequencing libraries. (b) Histogram of the number of microbial species per sample. (c) Bar chart of the human body sites that the 117 confidently detected species are associated with, as determined using the Disbiome database^34^. Species are classified as ‘multiple’ if they are associated with more than one body site and classified otherwise if they are only associated with a single body site. (d) Piechart showing the microbiological classification of the 117 confidently detected species. (e) Bar chart showing prevalence of genera in blood culture records and in the blood sequencing libraries before and after decontamination.

Given past reports of bacterial translocation from the mouth^32^ or gut^33^ into blood, we asked if the microbes we detected could have originated from various body sites. To do so, we assigned potential body site origins to the 117 microbial species detected in blood based on microbe-to-body-site mappings extracted from the Disbiome database^34^. We found that many (*n*=59; 50%) of these confidently detected species are indeed human commensals that are present at various human body sites (**Figure 2c**). While some of these species may be contaminants that have survived our stringent decontamination filters, this observation, together with their low prevalence, suggests that the microbial DNA of many of these species may have transiently translocated from other locations in the body rather than being endogenous to blood. We further categorised the microbial species based on their growth environments (**Figure 2d**). A significant portion (*n*=42; 36%) of the species were obligate anaerobes or obligate intracellular microbes, atypical of skin-associated microbes that may be introduced during phlebotomy^2^, indicating that they are not likely to be sampling artefacts. All in all, the diverse origins of the microbes detected in blood, together with their low prevalence across a healthy population, is consistent with sporadic translocation of commensals, or their DNA, into the bloodstream.

Microbial presence in blood (i.e., bacteraemia) is typically associated with a range of clinical sequelae from mild fevers to sepsis. As such, we asked if the common microbes identified in patients with disease-associated bacteraemia are different from those detected in our cohorts of healthy individuals. To do so, we compared the prevalence of microbes detected in the sequenced blood samples against observations from 11 years of hospital blood culture records. The prevalence of microbial genera detected in the hospital blood culture records clearly differed from that in our sequenced blood samples, despite the overlap in detected taxa (**Figure 2e**). For example, while *Staphylococcus, Escherichia* and *Klebisiella* were the predominant genera identified in blood cultures, they were rarely detected in our blood sequencing libraries. We performed a similar comparison with a previous study^35^ which sequenced blood microbial signatures in sepsis patients and found a similar difference in prevalence compared to our dataset (**Supplementary Figure 5**), confirming that our observations are not due to differences in the detection methods (sequencing vs. culture-based) used. If the species detected through sequencing were genuine, and represent microbial cells, these findings may be explained by the potentially higher virulence of pathogens detected in the clinic, which are more likely to cause clinical symptoms in individuals that would result in exclusion during our recruitment process. Conversely, under the same assumptions, our findings suggest that the microbes detected in the blood of healthy individuals are potentially better tolerated by the immune system (e.g. *Bifidobacterium* spp.^36^ and *Faecalibacterium prausnitizii*^37^ with immunomodulatory properties as gut commensals; **Figure 2a**).

### Evidence for replicating microbial cells but without community structure or host associations

To better characterise the microbial DNA signatures detected in blood, we asked if they reflect the presence of viable microbial cells as opposed to circulating cell-free DNA. This is because the former would allow for complex microbe-microbe or microbe-host interactions that would be of greater and more direct clinical relevance. In contrast to previous approaches that used microbial cultures^3,38^, we looked for more broad-based evidence of live bacterial growth in by applying replication rate analyses^21,22^ on our sequenced blood samples. This approach is based on the principle that DNA sequencing of replicating bacteria would yield an increased read coverage (i.e., peak) nearer to the origin of replication (*Ori*) and decreased coverage (i.e., trough) nearer to the terminus (*Ter*)^22^. A coverage peak-to-trough ratio (PTR) greater than one is indicative of bacterial replication. Through this analysis, we found evidence for replication of 11 bacterial species out of the 20 that were sufficiently abundant to do this analysis (**Figure 3a)**. The median-smoothed coverage plots of the replicating species all exhibited the sinusoidal coverage pattern (in black; **Figure 3b**) characteristic of replicating bacterial cells^22^. This contrasts with the even coverage patterns of three representative contaminants identified during the decontamination steps: *Achromobacter xylosoxidans, Pseudomonas mendocina* and *Alcaligenes faecalis* (**Figure 3c**). The *Ori* and *Ter* positions determined using coverage biases largely corresponded with an orthogonal method based on the GC-skew^39^ of bacterial genomes, suggesting that the replication rate analyses are reliable. Additionally, all but one of these replicating species are present in hospital blood culture records and in previous reports of bacteraemia^40–49^ (**Figure 3a**), indicating their ability to replicate in human blood. Overall, beyond the detection of microbial DNA, we present the first culture-independent molecular signatures for microbial replication from blood.

**Figure 3:**
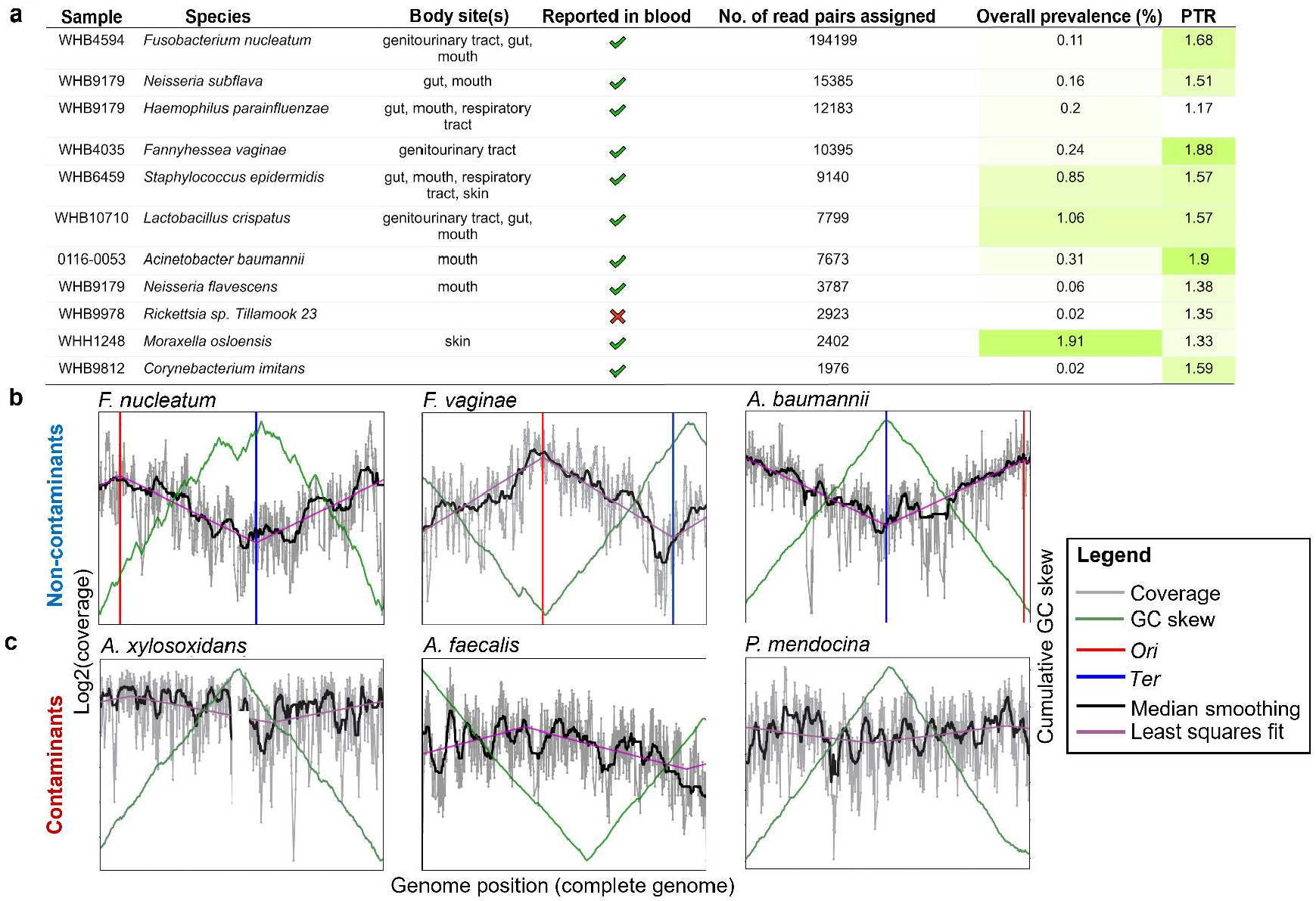
Evidence for replicating bacteria in blood samples from healthy individuals. (a) Summary statistics for samples where bacterial species were deemed to be replicating using *iRep*^21^ (i.e., peak-to-trough ratio (PTR)>1). The number of reads assigned to the species by *Kraken2* ^23^, the possible body sites the species are associated with, whether they were previously reported in published studies of bacteraemia, the overall prevalence of the species across all 8,892 individuals in our study and the calculated PTR values, are indicated. Coverage plots of (b) three representative confidently detected species and (c) three representative contaminant species, showing the expected patterns of *Ori* to *Ter* coverage skew only where expected i.e. confidently detected species.

Given the presence of live bacteria, we investigated if the microbial species detected showed patterns of microbe-microbe interactions as would be expected from a microbial community. To do so, we computed pairwise *SparCC* correlations^50^ between species, where positive and negative values indicate co-occurrence and mutual-exclusion, respectively. *SparCC* correlation is a reliable metric for assessing co-occurrence since it accounts for the sparse and compositional nature of microbial taxonomic profiles that confound standard correlation inference techniques^50^. We visualised *SparCC* correlations of the 117 microbial species confidently detected in blood sequencing libraries using network graphs, where each node is a species and each edge represents the co-occurrence/exclusion associations between two species (**Figure 4a**). We could not detect strong community co-occurrence/exclusion patterns, with most associations being weak (SparCC correlation<0.05), and only 19 pairwise associations exceeding a correlation value of 0.2, with four exceeding a value of 0.3 (**Figure 4a**). To determine if this result is a function of our stringent decontamination filters, we generated independent network graphs for the five adult cohorts before decontamination filtering and examined the co-occurrence/exclusion associations shared across cohorts. With an already lenient SparCC correlation threshold of 0.2, we identified no associations common to all the network graphs (**Figure 4b**), indicating that there were no consistent detectable microbial community associations in blood typical of microbiomes in various human body sites.

**Figure 4:**
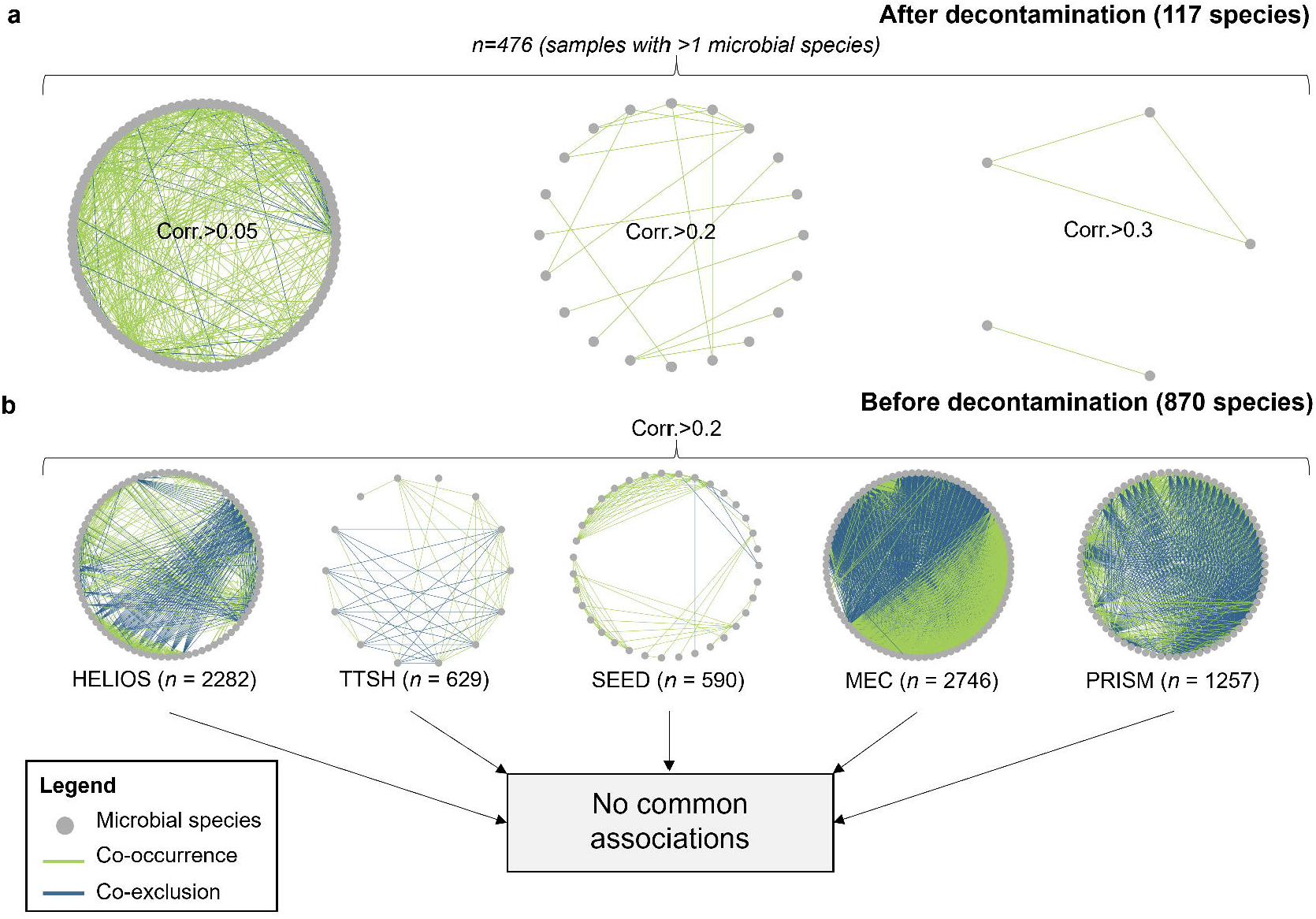
Microbial co-occurrence networks. (a) *SparCC*^50^ co-occurrence networks computed from all samples with at least two microbial species following decontamination at different *SparCC* correlation thresholds (0.05, 0.2, 0.3). Only associations with a magnitude of *SparCC* correlation greater than the respective thresholds are retained. (b) *SparCC* networks for individual cohorts at a correlation threshold of 0.2. No co-occurrence associations were retained after taking the intersection of edges between all cohort networks. For (a) and (b), each node represents a single microbial species, and each edge a single association between a pair of microbial species. Edge thickness is scaled by the magnitude of correlation. The number of samples used to compute each network and the correlation thresholds used are annotated. Positive and negative *SparCC* correlations are indicated in green and blue respectively.

Previous studies have demonstrated the use of blood microbial DNA as a biomarker for disease, demonstrating associations with cancer^30^, type II diabetes^51^ and periodontal disease^52^. In a similar vein, we investigated if the presence of microbes was associated with host phenotypes in our dataset. We first examined if microbes were detected more frequently in infants relative to adults. Given that the still-developing immune systems of infants puts them at greater risk of infection relative to healthy adults^53^, we reasoned that the prevalence of microbes in blood may differ within a birth cohort (GUSTO) relative to adult cohorts. Indeed, samples from GUSTO appeared to have a higher prevalence of microbes associated with most human body sites (**Supplementary Figure 6a**). This was in part, driven by genitourinary tract-associated microbes, *Fannyhessea vaginae, Lactobacillus jensenii, Lactobacillus crispatus, Lactobacillus iners*, and *Gardnerella vaginalis* (**Supplementary Figure 6b**). Similarly, we found enrichment of gut-associated bacteria such as *Bifidobacterium* spp. in GUSTO (**Supplementary Figure 6c**). These findings suggest that bacterial translocation may be more frequent in infants relative to adults, though differences in sample collection (umbilical cord *versus* venipuncture) could also explain them. A future study controlling for differences in sampling methods would be useful for further exploration of this observation.

Next, we systematically tested for pairwise associations between eight host phenotypes that were documented on the day of blood collection and the presence of each of the 117 microbial species detected in blood. These host phenotypes attributes were: sex, ancestry, age, body mass index (BMI), blood total cholesterol (TC), blood triglycerides (TG), systolic and diastolic blood pressure (SBP and DBP). Given the multiple large independent cohorts, we could perform statistical tests on each cohort separately, which allowed us to assess the consistency of identifed association patterns across the different cohorts. Since these cohorts were sampled from a homogenous population, true association patterns are expected to be detected repeatedly regardless of cohort. Using this statistical testing approach, we found only five significant microbe-phenotype associations (p<0.05; **Supplementary Table 3**) after adjusting for multiple comparisons. Notably, all but one of the significant associations were present in only one cohort. The exception was *C. acnes*, which was significantly associated with ancestry in two cohorts. However, while *C. acnes* was more prevalent in individuals of Malay ancestry within the SEED cohort, it was more prevalent in Chinese individuals within the MEC cohort (**Supplementary Figure 7**). These cohort specific differences could be due to other demographic variables that were not recorded in this study, or perhaps from *C. acnes* subspecies differences. To ensure that we did not miss any associations due to the possible non-linearity of host-phenotype and microbial relationships, we also derived categorical phenotypes based on the recorded phenotypic information. These include being elderly (age>=65), and other measures of ‘poorer health’, such as being obese (BMI>30), having high blood triglycerides (TG>2.3 mmol/L), high total cholesterol (TC>=6.3 mmol/L), or high blood pressure (SBP>=130 and DBP>=80). We then tested for pairwise associations between these derived phenotypes and the presence of *any* bacteria but found no significant associations (p>0.05; **Supplementary Table 4**). Collectively, these results suggest no consistent associations between the presence of microbes in blood and the host phenotypes tested within a healthy population of individuals.

## Discussion

We present the largest scale analysis, to date, of microbial signatures in human blood with rigorous accounting for computational and contamination artefacts and found no evidence for a common blood microbiome in a healthy population. Instead, we observed mostly sporadic instances of blood harbouring DNA from single microbial species of diverse bodily origins, some of which might be actively replicating. Our findings hint at the possibility that the bloodstream represents a route for movement of microbes between different body sites in healthy individuals. However, the low prevalence of the detected species suggest that this movement is likely to be infrequent and transient. Unresolved questions remain about how interconnected the microbiomes at various body sites are, and whether these processes are altered during disease or throughout a person’s lifetime. Can perturbations to the microbial community at one body site affect that at another site, and how does the host immune system asymptomatically regulate microbial presence in blood? Our study lays the groundwork for future investigations into these questions, which may pave the way for a systemic understanding of the human microbiome across body sites in relation to human health and disease.

We employed a series of decontamination filters to differentiate microbial signatures in blood from artefactual signals associated with reagent and handling contamination, on the basis that the latter display strong batch-specific biases (**Supplementary Figure 2;** see **Methods**). Although our approach substantially improved the signal-to-noise ratio (**Fig. 1b-d**), it is still likely not fully effective in removing contaminants, evidenced from the fact that 10% of the 117 microbial species remaining after decontamination were still flagged as being of environmental or non-human origin (**Fig. 1b**, “likely contaminant”). Hence, we recommend that any decontamination procedures should include further comparisons to various microbiome databases (**Fig. 1b-d**) to prioritise species for validation in future studies. For example, one might prioritise species that are not common contaminants, detected in blood cultures, and that are human associated (**Supplementary Table 2**) for follow-up experiments. Nevertheless, it is important to note that we could not detect a common blood microbiome despite the likely presence of residual contamination artefacts.

We observed signatures of replicating DNA from putatively genuine microbial species in blood by applying an existing PTR-based replication analysis approach. However, we cannot distinguish signals arising from replicating microbes in blood from those derived from microbial cells (intact or otherwise) which were recently replicating at other body sites before entering the bloodstream. Interestingly, while we could detect replication signatures in blood associated with 11 out of 20 species with sufficient coverage across their genomes, we could not detect any amongst the 20 most prevalent contaminant species identified by our decontamination filters, including species from the genera *Alcaligenes, Caulobacter, Bradyrhizobium* and *Sphingomonas*, suggesting that the replication signatures detected in our dataset are not likely to be due to ‘kitome’ contamination. Furthermore, this observation highlights the potential use of replication analyses for discriminating between putatively genuine taxa from ‘kitome’ contaminants in future metagenomic studies.

We found no core species in human blood on the basis of low prevalence across individuals in our population-level dataset. The prevalence estimates provided in this study are contingent on the sensitivity of detecting microbes through sequencing. Previous studies have shown that untargeted shotgun sequencing is highly sensitive for the detection of microbes in blood at a total sequencing depth of 20-30 million reads per sample^35,54,55^, perhaps even more so than culture-based methods^56,57^. In contrast, a median of 373 million reads was generated per sample for our sequencing libraries, suggesting that our methods do not lack sensitivity. Our prevalence estimates are also affected by the abundance thresholds used to determine whether a species is present in a single sample (i.e., abundance filter; **Figure 1a**). We defined these thresholds in terms of both absolute read count and relative abundance, which were determined based on simulation experiments (see **Methods**). Overly stringent abundance thresholds would lead to the erroneous masking of genuine signals, leading to an underestimation of microbial prevalence. However, even when relaxing the threshold to just a relative abundance of 0.001, none of the species, whether flagged as a contaminant or not, had more than 52% prevalence (**Supplementary Table 5**). Furthermore, the 20 most prevalent species at this threshold are all environmental microbes, and mostly comprise of *Sphingomonas* and *Bradyrhizobium* species, which are known to be common sequencing-associated contaminants^19^. This suggests that independent of our decontamination filters, none of the species detected qualify as core members.

In addition to not being able to detect any core species, we could not detect any strong co-occurrence or mutual exclusion associations between species regardless of whether our decontamination filters were applied. These associations generally reflect cooperation or competition between species, respectively^58^. Indeed, within a microbial community, metabolic dependencies of species and the ability of different species to complement these dependencies have been shown to be a key driver of microbial co-occurrence^59^. On the other hand, competitive behaviours such as nutrient sequestration to deprive potential competitors of nutrients or producing adhesins to bind and occupy favourable sites in an environment^60^ can lead to mutual exclusion between species. The fact that we could not detect any strong associations therefore points to the absence of an interacting microbial community in healthy humans. Of note, since our dataset was derived from circulating venous blood, we are, in principle, not able to detect microbial interactions that may be occurring at other sites of the bloodstream such as the inner endothelial lining of blood vessels. Experiments investigating the adherence of bacteria to blood vessel linings may provide further insight into this.

The availability of 11 years of blood culture records from the same country of origin as our blood samples enabled a reliable comparison of the prevalence of microbes in the healthy population and in the clinic. This is because the frequency of infections caused by different microbial species is known to differ from country to country^61^. Despite this, we expect that some of the variation in prevalence estimates may be due to the differences in detection methods. That said, previous studies have shown a strong concordance between culture and sequencing-based detection^35,54,56,57^, suggesting that the distinction between the prevalence of microbes found in healthy individuals and in the clinic is not due to the differences in detection methods. Our results support the conclusion that microbial presence in blood (i.e., bacteraemia) does not always lead to disease. These results are consistent with our other observation that microbial DNA detected in our cohorts of asymptomatic individuals tend to be from commensals, which may inherently be less virulent and better tolerated by the host compared to disease-causing pathogens. Indeed, the long-standing co-evolution of humans and colonizing microbes, places a selective pressure against high virulence phenotypes in these microbes to maintain host viability^62^. Simultaneously, there is a selective pressure for immunomodulatory phenotypes in commensals to improve their fitness, evidenced by the wealth of immunomodulatory activities found in the gut microbiome^63^. This agrees with previous findings that colonisation by commensals modulate early development of the immune system^64^, which would allow a measured and regulated response against translocated commensals. By extension, the immunomodulatory properties of bacteria and their links to host tolerance to bacteraemia may be key factors in determining clinical outcomes. Perhaps, the presence (or lack) of these properties may determine whether an individual with bacteraemia is asymptomatic or septic. For example, abundant gut bacterial species such as *Bacteroides* spp. were not commonly detected in blood. Further exploration into the immunomodulatory activities of commensals vis-à-vis common blood culture pathogens may be the key to design therapeutics to manage or prevent the dysregulated host response that defines sepsis^1^.

We found no convincing associations between both measured (e.g. TC, SBP) and derived (e.g. obesity) host phenotypes with microbial presence that were consistent across the different cohorts. This suggests that the risk of transient microbial translocation, at least across our cohorts of healthy adults, is fairly consistent. In contrast, this risk may increase in individuals with more severe disease. In fact, variable microbial DNA profiles in blood have been used to delineate health and disease states. This has most prominently been shown for sepsis^35,54–57,65^, where the presence of viable microbes is expected, but also for cancer^30^, periodontal disease^52^, and chronic kidney disease^66^, which are unrelated to bloodstream infections. These studies highlight the promise of metagenomic sequencing of blood for developing diagnostic, prognostic, or therapeutic tools. Our characterisation of the species breadth in healthy individuals forms a crucial baseline for comparison with that in diseased individuals. Indeed, our findings open new doors to understanding why and how blood microbial profiles correlate with health status. One possible hypothesis is that mucosal integrity is compromised in a disease state, leading to higher translocation rates of microbes into the bloodstream. This is consistent with findings of increased intestinal permeability (i.e., ‘leaky gut’) in disease or even during physiological stress^67^. Future studies testing this hypothesis may consider a focus on the gut-associated bacteria that were detected in our study (e.g. *Bifidobacterium adolescentis, Faecalibacterium prausnitzii*). Further experimental investigations into the mechanisms of microbial translocation and the modulatory effects of the microbiomes present at other body sites may shed light on the relationship between microbial presence in blood and health status.

If we take the definition of a ‘microbiome’ as a microbial community whose member species interact amongst themselves and with their ecological niche^9^, our findings lead to the conclusion that there is no consistent circulating blood microbiome in healthy individuals. Sporadic and transient translocation of commensals from other body sites into the bloodstream (**Figure 5**) is the more parsimonious explanation for the observation that most of the microbes detected are commensals from other body sites. Furthermore, the relatively low prevalence of microbes in blood suggests rapid clearance of translocated microbes rather than prolonged colonisation in blood. Based on these findings, we advocate against the use of the term ‘blood microbiome’ or ‘circulating microbiome’, which are potentially misleading, when referring to the detection of microbial DNA or of microbial cells in blood due to transient translocation events.

**Figure 5:**
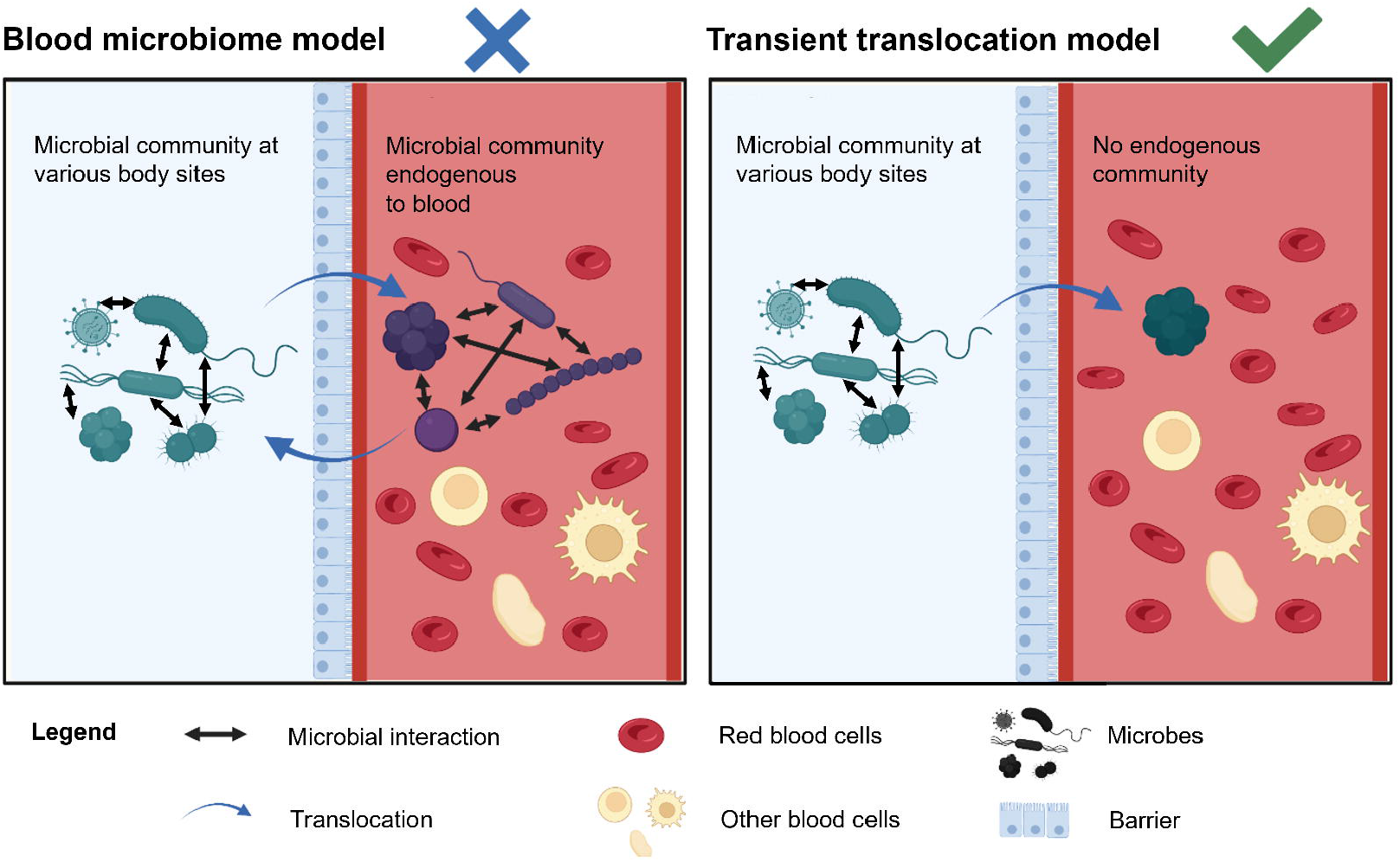
Potential models for microbes in blood. Our findings suggest that there is no consistent circulating blood microbiome (i.e., the blood microbiome model). The more likely model is where microbes from other body sites transiently and sporadically translocate into blood. Created with BioRender.com under an academic subscription.

## Methods

### Datasets

Our sequencing dataset, also known as the SG10K_Health dataset (https://www.npm.sg/collaborate/partners/sg10k/), comprises of shotgun sequencing libraries of DNA extracted from the whole blood or umbilical cord blood of 9,770 healthy Singaporean individuals^68^ who were recruited as part of six independent cohorts. Individuals were deemed to be healthy if they do not have any personal history of major disorders such as stroke, cardiovascular diseases, cancer, diabetes and renal failure. Oral health information was not collected and therefore not part of the exclusion criteria. Whole blood for sequencing was collected via venipuncture only from the five adult cohorts (median age=49; interquartile range=16): Health for Life in Singapore (HELIOS; *n*=2,286), SingHealth Duke-NUS Institute of Precision Medicine (PRISM, *n*=1,257), Tan Tock Seng Hospital Personalised Medicine Normal Controls (TTSH, *n*=920), Singapore Epidemiology of Eye Diseases (SEED, *n*=1,436)^69,70^, and the Multi-Ethnic Cohort (MEC, *n*=2,902)^71^. Additionally, cord blood was collected only for the birth cohort Growing Up in Singapore Towards healthy Outcomes (GUSTO; *n*=969)^72^. Measurement of host phenotypes was performed on the day of blood collection, except for the GUSTO cohort where measurements were taken at a later timepoint when the children were at a median age of 6.1 (interquartile range=0.1). Using nearest neighbor approaches to reference genotypes^73^, individuals were broadly categorised into four ethnic categories representing distinct genetic ancestries: Chinese (59%), Malays (19%), Indians (21%) and Others (1%). All individuals were deemed healthy at the point of recruitment if they did not include any self-reported diseases in the recruitment questionnaires. All cohort studies were approved by relevant institutional ethics review boards. A summary of the cohort demographics and the ethics review approval reference numbers are provided in **Supplementary Table 1**.

Additionally, we retrieved anonymised blood culture records from Singapore General Hospital, the largest tertiary hospital in Singapore. These records span the years 2011-2021 and include aerobic, anaerobic and fungal blood cultures taken from 282,576 unique patients. These blood cultures were ordered as part of routine clinical management, that is, when clinically indicated for the investigation of bacteremia or fungemia. Blood cultures were performed and analysed as per hospital standard operating procedures. In brief, blood samples were collected aseptically and inoculated into BD™ BACTEC™ bottles at the bedside (BD™ BACTEC™ Plus Aerobic/F Culture vials Plastic [catalogue number 442023] for aerobic blood culture, BD™ BACTEC™ Plus Anaerobic/F Culture vials Plastic [catalogue number 442022] for anaerobic blood culture and Myco/F Lytic [catalogue number 42288] for fungal blood culture). The inoculated bottles were transported to the diagnostic laboratory at ambient temperature and incubated in the BD™ BACTEC™ FX Blood Culture System on arrival. Aerobic and anaerobic blood culture bottles were incubated for a maximum of five days, and fungal blood culture bottles were incubated for a maximum of 28 days. Blood culture bottles that were flagged positive by the BD™ BACTEC™ FX Blood Culture System were inoculated onto solid media, and the resultant colonies were identified using a combination of biochemical tests and matrix assisted laser desorption ionization-time of flight mass spectrometry (MALDI-TOF MS) (Bruker^®^ microflex LRF).

### Sample preparation and batch metadata

DNA from whole blood was extracted using one of six different DNA extraction kits. Paired-end 151bp sequencing with an insert size of 350bp was performed up to 15-fold or 30-fold coverage of the human genome. Library preparation was performed using one of three library preparation kits. Sequencing was performed on the Illumina HiSeq X platform with HiSeq PE Cluster Kits and HiSeq SBS Kits. The type of extraction kits and library preparation kits used, and lot numbers for the SBS Kits, PE Cluster Kits, and sequencing flow cells used are provided as batch metadata. All reagent kits used, the number of batches and the number of samples processed per batch are provided in **Supplementary Table 6**.

### Data pre-processing and quality control

The bioinformatic processing steps applied to the sequencing libraries are summarised in **Figure 1a**. Read alignment of sequencing reads to the GRCh38 human reference genome was already performed as part of a separate study^68^ using *BWA-MEM v0*.*7*.*17*^74^. We retrieved read pairs where both members of the pair did not map to the human genome. Following which, we performed quality control of the sequencing reads. We trimmed low quality bases at the ends of reads with quality <Q10 (base quality trimming) and discarded reads with average read quality less than Q10 (read quality filter). We also discarded low complexity sequences with an average entropy less than 0.6, with a sliding window of 50 and k-mer length of five (low complexity read filter). All basic quality control steps were performed using *bbduk* from the *BBTools suite* v37.62 (sourceforge.net/projects/bbmap/).

### Taxonomic classification of blood sequencing libraries

Taxonomic classification of non-human reads was done using *Kraken2* v2.1.2^23^ with the ‘—paired’ flag. We used the *PlusPF* database (17^th^ May 2021 release) maintained by Ben Langmead (https://genome-idx.s3.amazonaws.com/kraken/k2_pluspf_20210517.tar.gz), which includes archaeal, bacterial, viral, protozoan, and fungal references. Of all non-human read pairs, 72% were classified as microbial at the species level, yielding 8,890 species. Samples with less than 100 microbial read pairs were removed, resulting in a final dataset comprising 8,892 samples, with a median microbial read-pair count of 6187.

To minimise noise in the taxonomic assignments, we defined a set of abundance thresholds whereby species with abundance values less than or equal to these thresholds (i.e., relative abundance≤0.005, read pairs assigned≤10) were counted as absent (set to zero read counts). We performed simulations to systematically determine a relative abundance threshold that minimizes false positive species assignments. Sequencing reads were simulated using *InSilicoSeq v1*.*5*.*4*^75^ with error models trained on the SG10K_Health sequencing libraries and processed using the same bioinformatic steps as per the SG10K_Health dataset to obtain microbial taxonomic profiles. We simulated 373 million reads equivalent to the median library read count of all samples, comprising reads from the GRCh38 human reference and ten microbial genomes (*Yersinia enterocolitica, Leclercia adecarboxylata, Moraxella osloensis, Streptococcus pneumoniae, Pasteurella multocida, Staphylococcus epidermidis, Actinomyces viscosus, Torque teno virus, Human betaherpesvirus 6A, Candida albicans*) at various proportions. Due to read misclassification, some of the simulated reads were erroneously assigned to another species and produced false positives. A final relative abundance threshold of 0.005 that delineated these false positive assignments from true positives was selected (**Supplementary Figure 8**). Following the application of these thresholds, the relative abundance distribution of microbial taxa classified as present were distinct from the distribution for those classified as absent (**Supplementary Figure 9**). Furthermore, the distribution of abundances for microbe-negative samples is centred around a relative abundance of 0.0001, i.e. at least tenfold below the typical relative abundance thresholds used to determine if a taxon is present or absent (0.001-0.045^14^). Relative abundances were calculated by dividing the microbial read count in a sample by the total number of microbial reads assigned to that sample.

### Decontamination filters

After application of the presence/absence filter, we identified and removed putative contaminants using established decontamination heuristics^26^ that have been validated in previous studies^27,28^, prior to our downstream analyses. These rules were applied using eight types of batch information: source cohort, DNA extraction kit type, library preparation kit type, and lot numbers for sequencing-by-synthesis kit (box 1, box 2), paired-end cluster kit (box 1, box 2) and sequencing flow cell used. Other batch information such as the pipettes and consumables used, or storage location and duration were not recorded and could potentially contribute to some level of batch-specific contamination. However, these batches are expected to be correlated with the other types of batch information available, and so the resultant contaminants could in theory be accounted for using our filters. We describe the four decontamination filters used, as shown in **Figure 1a**, in sequential order:

1. *Prevalence filter*. A microbial species is considered a contaminant specific to a batch if it is present at greater than 25% prevalence in that batch and has greater than a two-fold higher prevalence than that for any other batch. Batches with less than 100 samples were excluded from this analysis. This filter is based on the principle that species which are highly prevalent in some batches but lowly prevalent or absent in others are likely contaminants^26^. We illustrate this for an example species in **Supplementary Figure 10a**.
2. *Correlation filter*. A microbial species is considered a contaminant if it is highly correlated (Spearman’s ρ>0.7) with any contaminant within the same batch, as identified by the prevalence filter. This filter is based on the principle that contaminants are highly correlated within the same batch^26^. Spearman’s ρ was calculated using centred log-ratio (CLR) transformed^76^ microbial relative abundances. CLR transformations and Spearman’s ρ were calculated using the *clr* function as part of the *compositions* package^77^ and *cor*.*test* function in *R*. We illustrate this within-batch correlation for an example species in **Supplementary Figure 10b**.
3. *Batch filter*. A non-contaminant microbial species must be detected in samples processed by at least two reagent kit batches or reagent types. That is, any species that is only detected in a single batch for any of the reagent kits used (**Supplementary Table 6**) are considered contaminants. This filter is based on the principle that species that can be repeatedly observed across different reagent batches are more likely to reflect genuine non-contaminant signals^26^. Library preparation kit type was excluded from this analysis since only three kit types were used, with 86% of samples processed using one of the kits.
4. *Read count filter*. A microbial species is considered a sequencing or analysis artefact if it is not assigned at least 100 reads in at least one sample. This filter is based on the principle that species that are always assigned a low number of read pairs, never exceeding the background noise within sequencing libraries, are more likely to be artefactual rather than genuine signals. An example of an artefactual species is *Candidatus Nitrosocosmicus franklandus*, which was assigned at most 22 read pairs by *Kraken2* across 21 sequenced samples.

To demonstrate the effectiveness of our decontamination filters, we additionally tested our results against the null hypothesis that the 117 microbial species retained after decontamination produced the same proportions of species classified as likely contaminants, human-associated, or that were detected in blood culture compared to if we picked these species at random. In this analysis, we generated 1000 sets of 117 microbial species that were randomly selected from the list of species before decontamination and compared the species to the three databases as per **Figure 1b-d**. P-values were calculated by taking the proportion of random iterations that generated proportions of species classified as likely contaminants, detected in blood, or human-associated that were as extreme or more extreme than those observed for the 117 species retained by our decontamination filters.

### Characterisation of microbial species

We classified microbial species as human-associated or not based on a published host-pathogen association database^78^. In this database, host-pathogen associations are defined by the presence of at least one documented infection of the host by the pathogen^31^. For species that were not found in this database, we performed a systematic PubMed search using the search terms: (microbial species name) AND (human) AND ((infection) OR (commensal)). Similarly, species that had at least one published report of human colonisation/infection were considered human-associated. Additionally, we classified the potential body site origins for each microbial species using the *Disbiome* database, which collects data and metadata of published microbiome studies in a standardised way^34^. We extracted the information for all microbiome experiments in the database using the URL: ‘https://disbiome.ugent.be:8080/experiment’ (accessed 26^th^ April 2022). We first extracted microbe-to-sample type mappings from this information (e.g. *C. acnes*→skin swab). We then manually classified each sample type into different body sites (e.g. skin swab→skin). This allowed us to generate microbe-to-body site mappings. Sample types with ambiguous body site origins (e.g. abscess pus) were excluded. The range of sample types within the Disbiome database used to derive the microbe-body-site mappings are provided in **Supplementary Table 7**. Finally, we classified microbial species based on their growth requirements, with reference to a clinical microbiology textbook^79^. Viruses were classified as obligate intracellular. The microbiological classifications for each species are provided in **Supplementary Table 2**.

### Estimating coverage breadth and bacterial replication rates

We performed read alignment of sequencing libraries to microbial reference genomes using *Bowtie v2*.*4*.*5*^80^ with default parameters. In total, we used references for 28 of the 117 microbial species detected in blood, comprising all bacterial species with at least 1000 Kraken2-assigned read pairs in a single sample and all viral species (*n*=5). For each species, we aligned the microbial reads of five sample libraries with the most reads assigned to that species, to the reference genome of that species. For each sample and microbial genome, the genome coverage per position was computed using the *pileup* function as part of the *Rsamtools v2*.*8*.*0* package^81^ in *R*. In principle, recovery of a larger fraction of a microbial genome provides a higher confidence that it is truly present in a sample^24,25^. We could recover at least 10% of the microbial genomes for 27/28 (96%) of the species. Since it is difficult to assess coverage breadth for a species covered by a low number of reads, we only performed this analysis on all viruses (*n*=5), and all bacterial species with at least 1000 Kraken2-assigned read pairs (*n*=23), which corresponds to ∼10% coverage over a typical 3Mbp bacterial genome (assuming non-overlapping reads). For the replication rate analyses, PTR values were calculated using the *bPTR* function in *iRep v1*.*1*.*0*^21^, which is based on the method proposed by Korem et al.^22^. The *Ori* and *Ter* positions were determined based on the coverage peaks and troughs (in red and blue, respectively; **Figure 3**). *Ori* and *Ter* positions were also calculated using a cumulative GC-skew line, which is expected to be in anti-phase with the sinusoidal coverage pattern across the genome^39^ (in green; **Figure 3**).

### Microbial networks

Microbial co-occurrence/mutual exclusion associations were computed using the *SparCC* algorithm^50^, implemented in the *SpiecEasi v1*.*1*.*2* package^82^ in *R* and the microbial networks were visualized using *Igraph v1*.*2*.*9*^83^. We excluded the birth cohort GUSTO since it is of a different demographic that may possess a distinct set of microbial associations.

### Detecting associations between microbial taxonomic profiles and host phenotypes

We tested for microbe-host phenotype associations within individual cohorts separately. For the two categorical host phenotypes, genetic sex and ancestry, we tested for differences in the prevalence of each microbial species between the different categories using a two-sided Fisher’s exact test (*fisher*.*test* function in *R*). For the continuous variables (age, BMI, TC, TG, SBP and DBP) we used a two-sided Mann-Whitney U test (*wilcox*.*test* function in *R*) to test for differences in the distributions of the variables when a species was present or absent. Benjamini-Hochberg multiple-testing correction was applied only after consolidating the *p*-values from both tests and for all cohorts using the *p*.*adjust* function in *R*. Statistical tests were only performed if a species was present in at least 50 samples in total. Separately, for derived phenotypes (i.e., being elderly or measures of ‘poorer health’), we used the Fisher’s exact test before applying Benjamini-Hochberg multiple-testing correction. In all cases, samples with missing host phenotypes were excluded.

### Data analysis and visualisation

All data analyses were performed using *R* v4.1.0 or using *Python* v3.9.12. Visualisations were performed using *ggplot v3*.*3*.*5*^84^. **Figure 5** was created using BioRender.com under an academic subscription.

## Data availability

Requests for the sequencing data used in this study should be made through the National

Precision Medicine (NPM) Programme Data Access Committee. All other data used in our analyses are hosted on Zenodo (https://doi.org/10.5281/zenodo.7368262). The accession numbers for all genome references used are provided in **Supplementary Table 8**.

## Code availability

All custom code used to perform the analyses reported here are hosted on GitHub (https://github.com/cednotsed/blood_microbial_signatures.git).

## Acknowledgements

We would like to thank the SG10K_Health Consortium, whose members and affliations are listed in **Supplementary Table 9**, for the collection and curation of the data used in this study. The computational work for this article was partially performed on resources of the National Supercomputing Centre, Singapore (https://www.nscc.sg). This study made use of data generated as part of the Singapore National Precision Medicine program funded by a grant from the Industry Alignment Fund (Pre-Positioning) (IAF-PP: H17/01/a0/007).

This study made use of data / samples collected in the following cohorts in Singapore:

1. The Health for Life in Singapore (HELIOS) study at the Lee Kong Chian School of Medicine, Nanyang Technological University, Singapore (supported by grants from a Strategic Initiative at Lee Kong Chian School of Medicine, the Singapore Ministry of Health (MOH) under its Singapore Translational Research Investigator Award (NMRC/STaR/0028/2017) and the IAF-PP: H18/01/a0/016);
2. The Growing up in Singapore Towards Healthy Outcomes (GUSTO) study, which is jointly hosted by the National University Hospital (NUH), KK Women’s and Children’s Hospital (KKH), the National University of Singapore (NUS) and the Singapore Institute for Clinical Sciences (SICS), Agency for Science Technology and Research (A*STAR) (supported by the Singapore National Research Foundation under its Translational and Clinical Research (TCR) Flagship Programme and administered by the Singapore Ministry of Health’s National Medical Research Council (NMRC), Singapore - NMRC/TCR/004-NUS/2008; NMRC/TCR/012-NUHS/2014. Additional funding was provided by SICS and IAF-PP H17/01/a0/005);
3. The Singapore Epidemiology of Eye Diseases (SEED) cohort at Singapore Eye Research Institute (SERI) (supported by NMRC/CIRG/1417/2015; NMRC/CIRG/1488/2018; NMRC/OFLCG/004/2018);
4. The Multi-Ethnic Cohort (MEC) cohort (supported by NMRC grant 0838/2004; BMRC grant 03/1/27/18/216; 05/1/21/19/425; 11/1/21/19/678, Ministry of Health, Singapore, National University of Singapore and National University Health System, Singapore);
5. The SingHealth Duke-NUS Institute of Precision Medicine (PRISM) cohort (supported by NMRC/CG/M006/2017_NHCS; NMRC/STaR/0011/2012, NMRC/STaR/0026/2015, Lee Foundation and Tanoto Foundation);
6. The TTSH Personalised Medicine Normal Controls (TTSH) cohort (supported by NMRC/CG12AUG17 and CGAug16M012).

The views expressed are those of the authors, are not necessarily those of the National Precision Medicine investigators, or institutional partners. We thank all investigators, staff members and study participants who made the National Precision Medicine Project possible.

## Ethics declaration

All individuals in the participating cohorts were recruited with signed informed consent from the participating individual or parent/guardian in the case of minors. All studies were approved by relevant institutional ethics review boards detailed in **Supplementary Table 1**.

## Notes

### Competing Interest Statement

The authors have declared no competing interest.

### Summary of Updates

Inclusion of new analysis and supplementary figures

