## Supplementary Figures 1-10 for "No evidence for a common blood microbiome based on a population study of 9,770 healthy humans"

### Slide 1
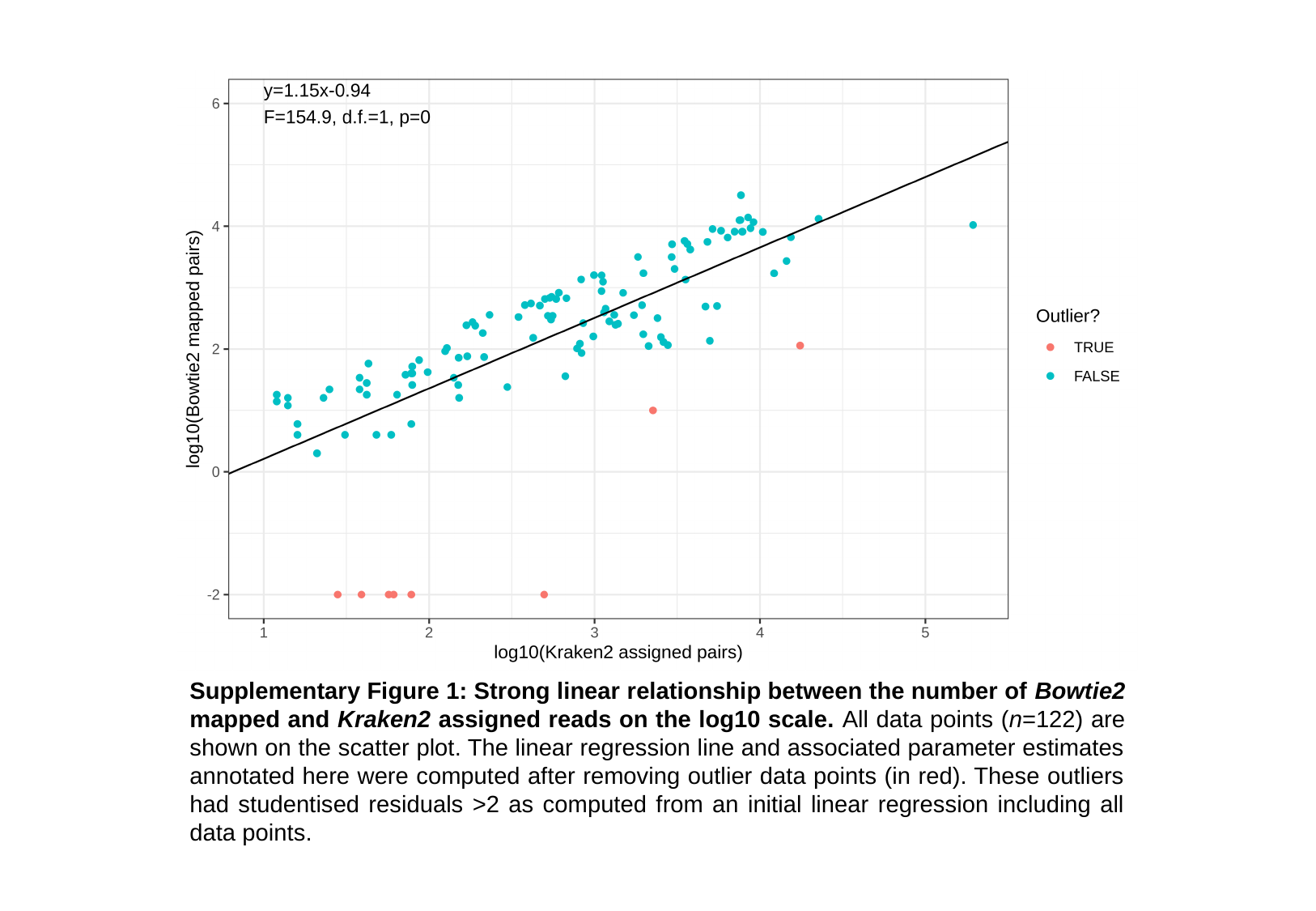

Supplementary Figure 1: Strong linear relationship between the number of Bowtie2 mapped and Kraken2 assigned reads on the log10 scale. All data points (n=122) are shown on the scatter plot. The linear regression line and associated parameter estimates annotated here were computed after removing outlier data points (in red). These outliers had studentised residuals >2 as computed from an initial linear regression including all data points.

### Slide 2
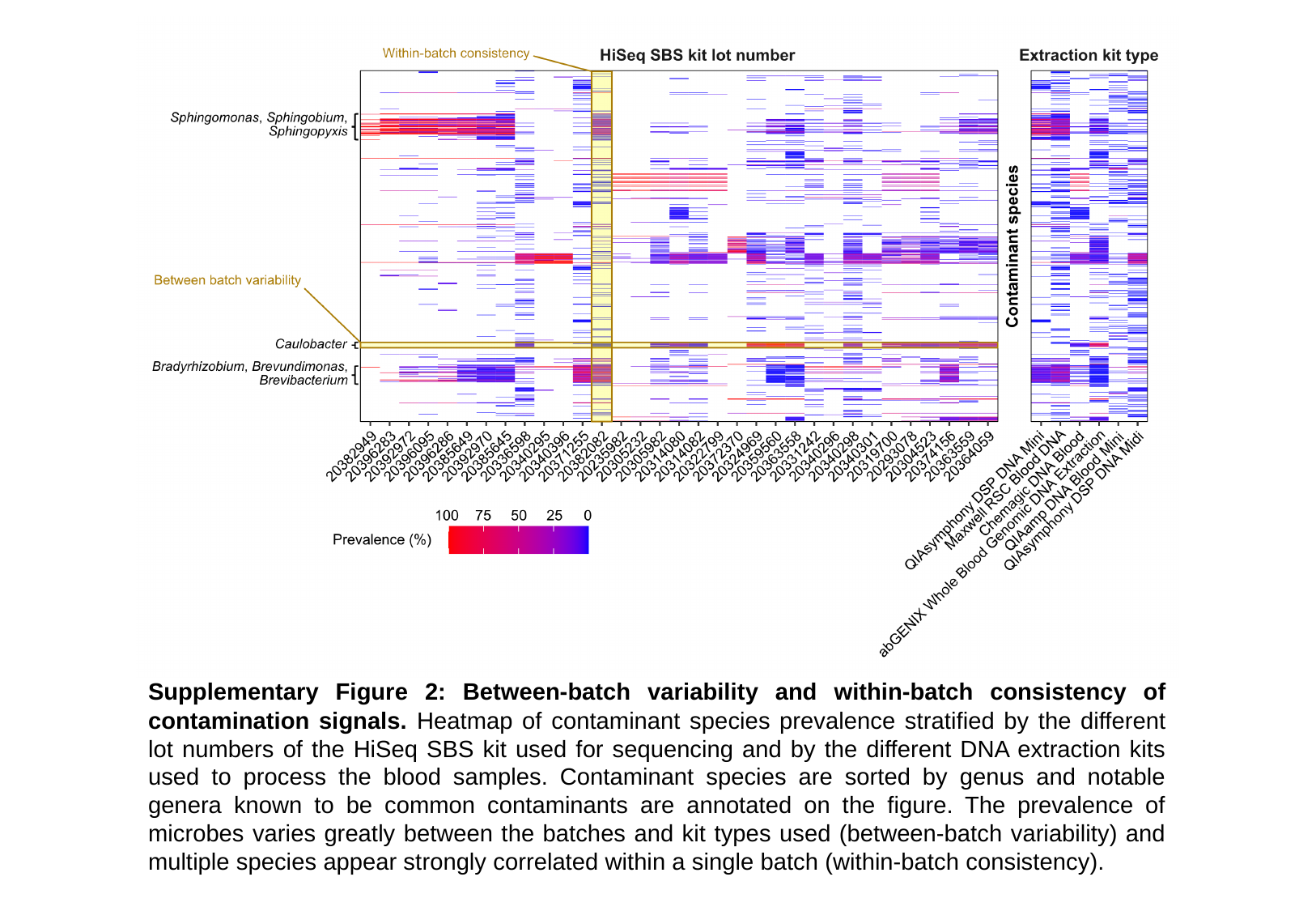

Supplementary Figure 2: Between-batch variability and within-batch consistency of contamination signals. Heatmap of contaminant species prevalence stratified by the different lot numbers of the HiSeq SBS kit used for sequencing and by the different DNA extraction kits used to process the blood samples. Contaminant species are sorted by genus and notable genera known to be common contaminants are annotated on the figure. The prevalence of microbes varies greatly between the batches and kit types used (between-batch variability) and multiple species appear strongly correlated within a single batch (within-batch consistency).

### Slide 3
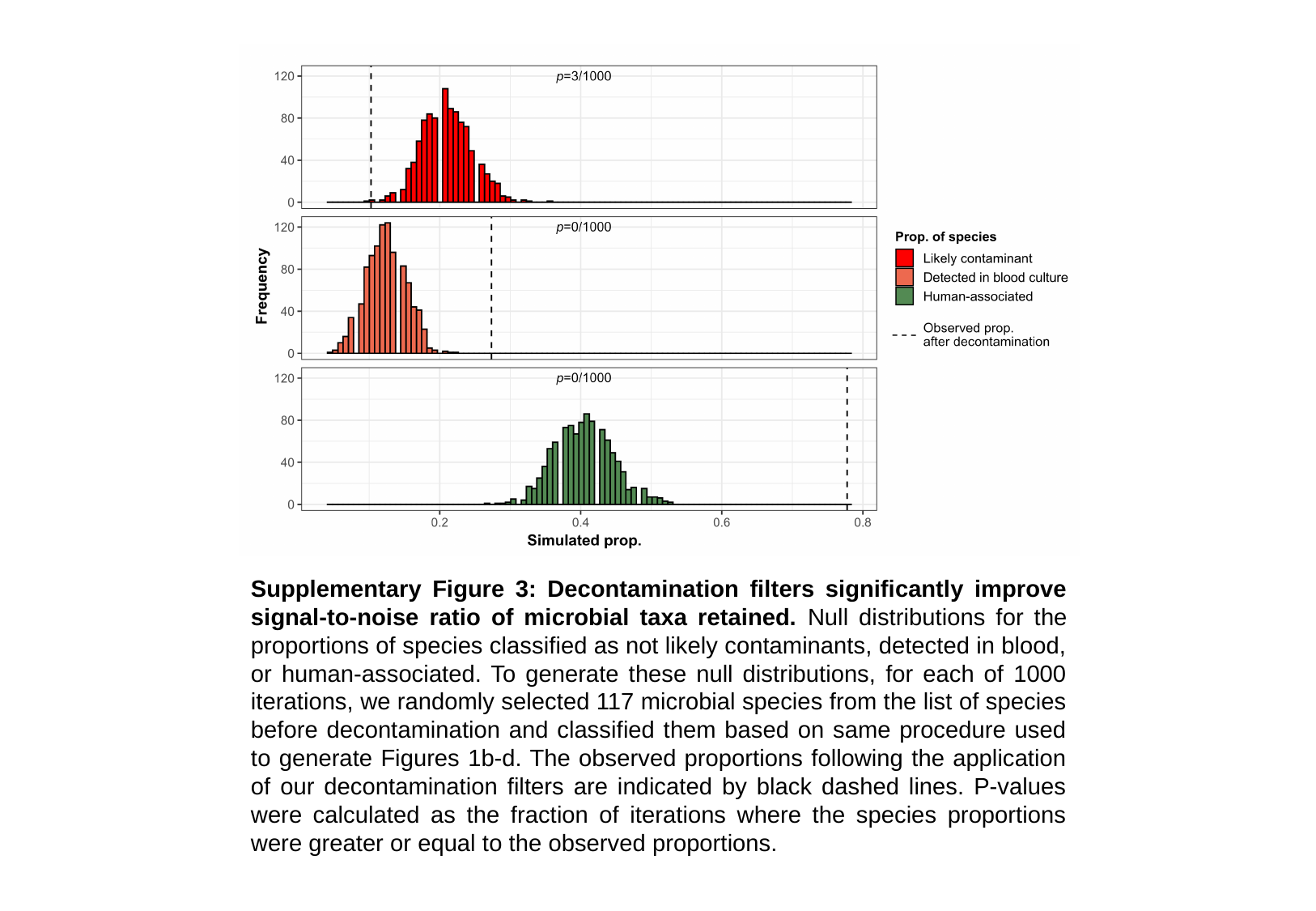

Supplementary Figure 3: Decontamination filters significantly improve signal-to-noise ratio of microbial taxa retained. Null distributions for the proportions of species classified as not likely contaminants, detected in blood, or human-associated. To generate these null distributions, for each of 1000 iterations, we randomly selected 117 microbial species from the list of species before decontamination and classified them based on same procedure used to generate Figures 1b-d. The observed proportions following the application of our decontamination filters are indicated by black dashed lines. P-values were calculated as the fraction of iterations where the species proportions were greater or equal to the observed proportions.

### Slide 4
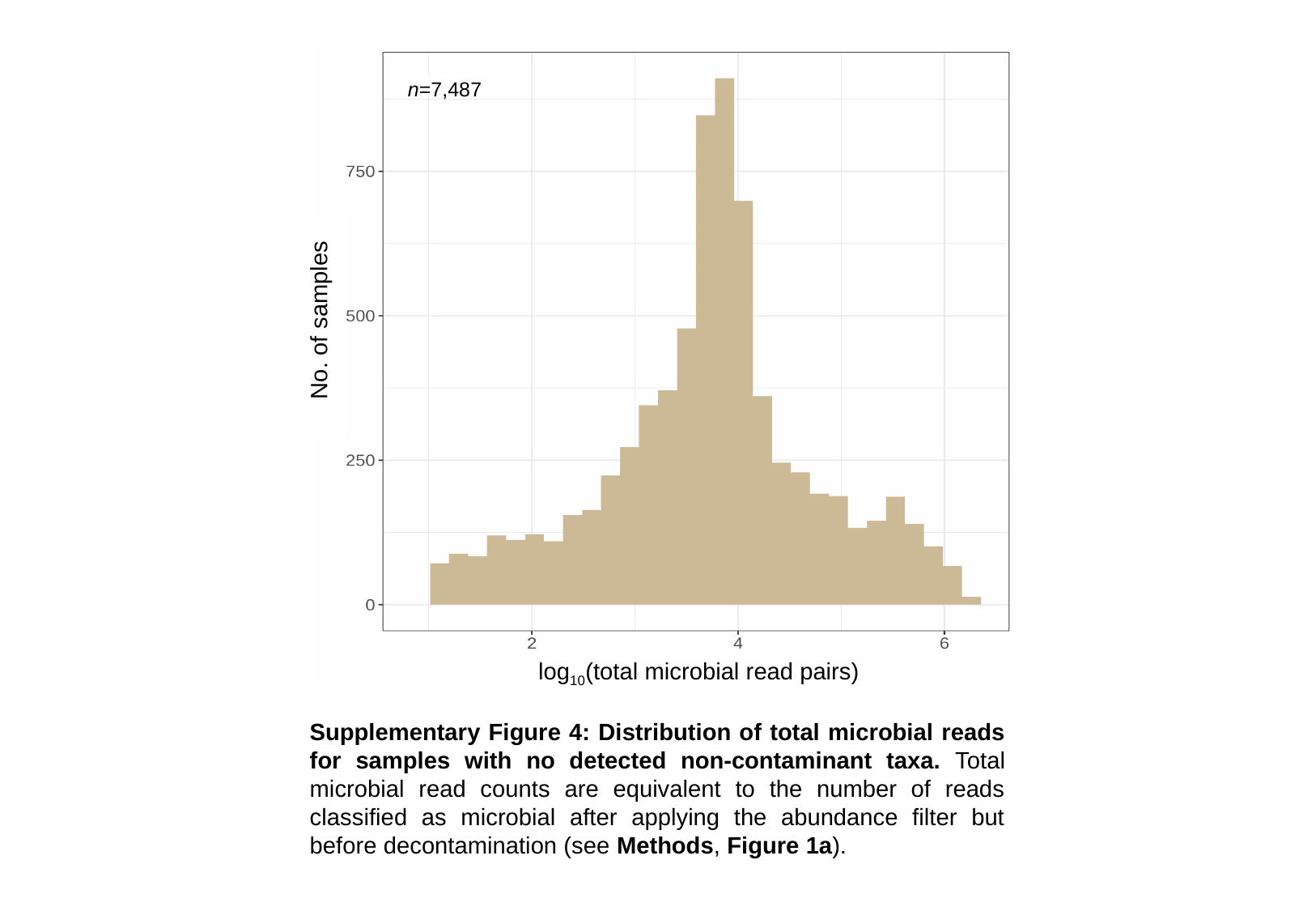

n=7,487
No. of samples
log_10_(total microbial read pairs)
Supplementary Figure 4: Distribution of total microbial reads for samples with no detected non-contaminant taxa. Total microbial read counts are equivalent to the number of reads classified as microbial after applying the abundance filter but before decontamination (see Methods, Figure 1a).

### Slide 5
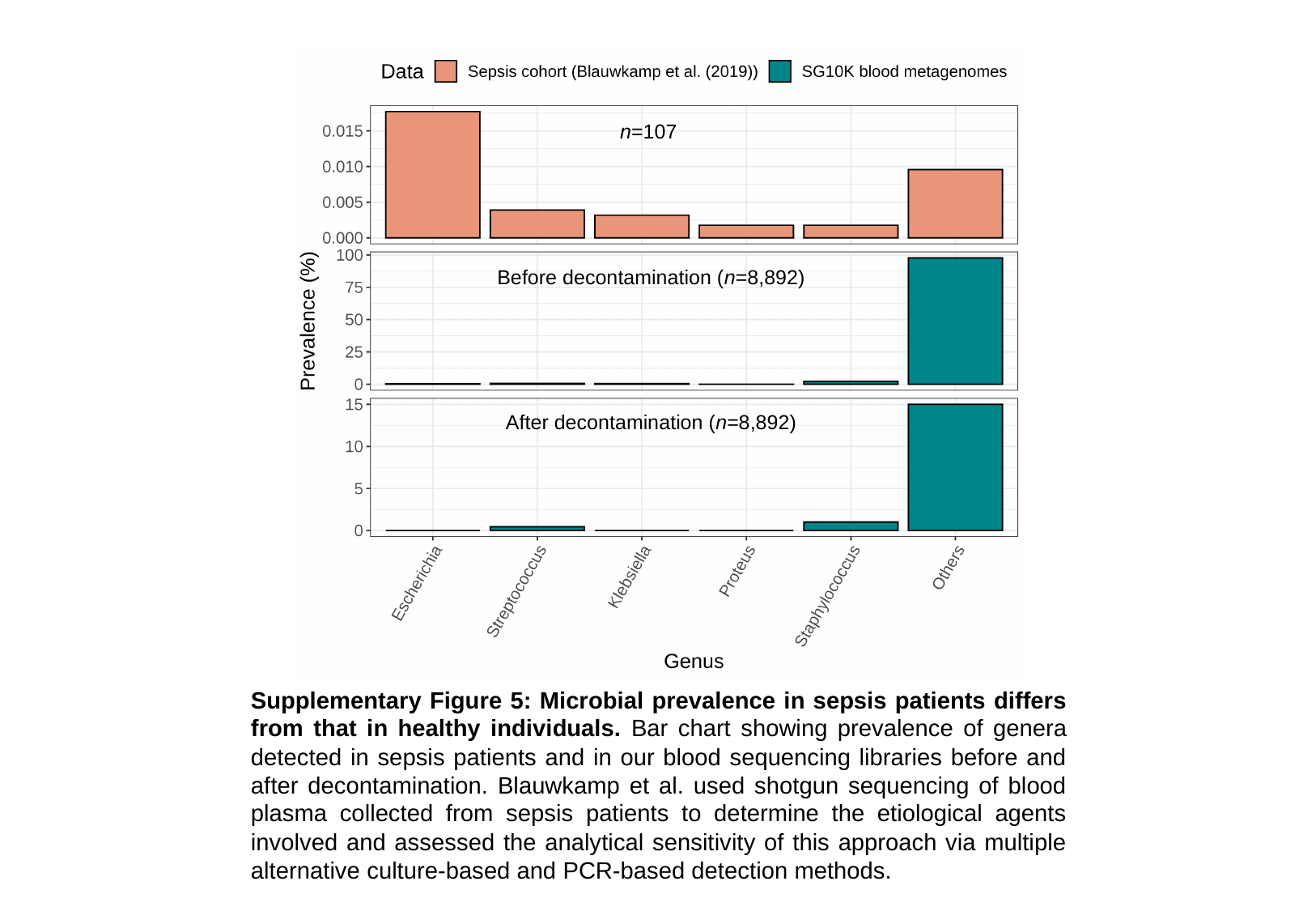

n=107
Before decontamination (n=8,892)
After decontamination (n=8,892)
Supplementary Figure 5: Microbial prevalence in sepsis patients differs from that in healthy individuals. Bar chart showing prevalence of genera detected in sepsis patients and in our blood sequencing libraries before and after decontamination. Blauwkamp et al. used shotgun sequencing of blood plasma collected from sepsis patients to determine the etiological agents involved and assessed the analytical sensitivity of this approach via multiple alternative culture-based and PCR-based detection methods.

### Slide 6
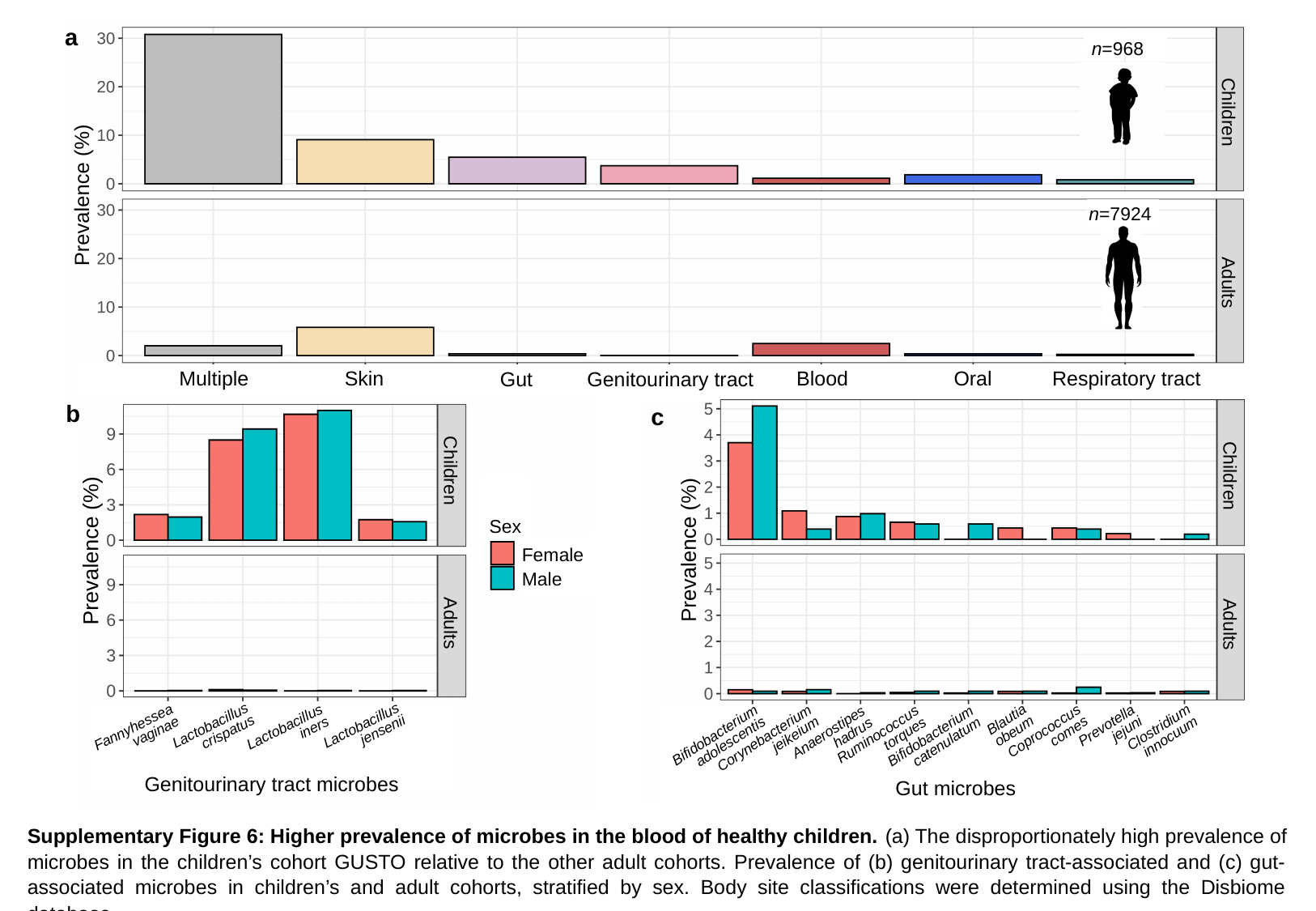

a
n=968
Children
n=7924
Adults
Oral
Multiple
Skin
Blood
Respiratory tract
Genitourinary tract
Gut
b
c
Children
Children
Sex
Female
Male
Adults
Adults
Blautia obeum
Lactobacillus crispatus
Lactobacillus jensenii
Fannyhessea vaginae
Lactobacillus iners
Prevotella jejuni
Coprococcus comes
Bifidobacterium adolescentis
Anaerostipes hadrus
Bifidobacterium catenulatum
Ruminococcus torques
Clostridium innocuum
Corynebacterium jeikeium
Genitourinary tract microbes
Gut microbes
Gut microbes
Supplementary Figure 6: Higher prevalence of microbes in the blood of healthy children. (a) The disproportionately high prevalence of microbes in the children’s cohort GUSTO relative to the other adult cohorts. Prevalence of (b) genitourinary tract-associated and (c) gut-associated microbes in children’s and adult cohorts, stratified by sex. Body site classifications were determined using the Disbiome database.

### Slide 7
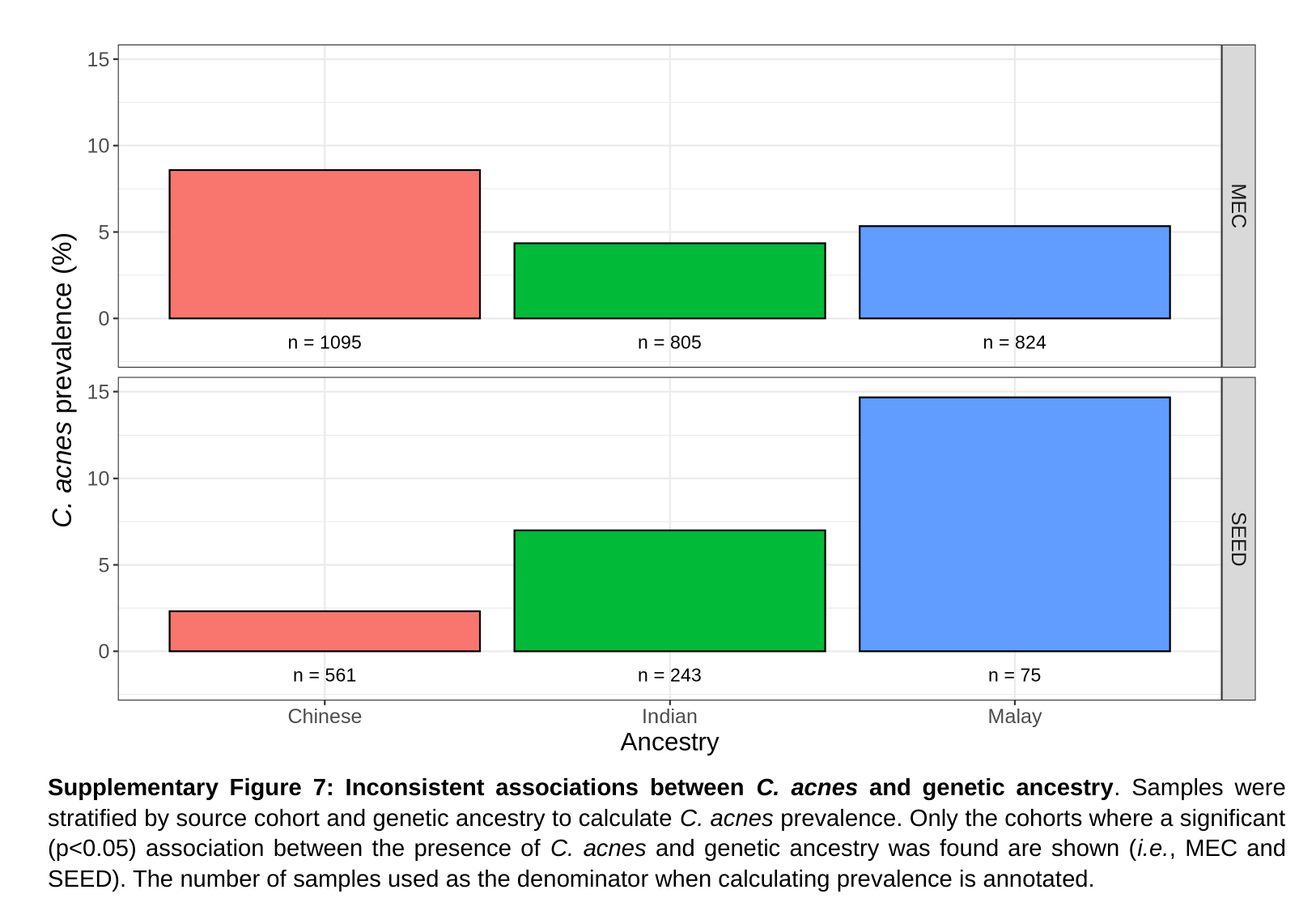

C. acnes prevalence (%)
Supplementary Figure 7: Inconsistent associations between C. acnes and genetic ancestry. Samples were stratified by source cohort and genetic ancestry to calculate C. acnes prevalence. Only the cohorts where a significant (p<0.05) association between the presence of C. acnes and genetic ancestry was found are shown (i.e., MEC and SEED). The number of samples used as the denominator when calculating prevalence is annotated.

### Slide 8
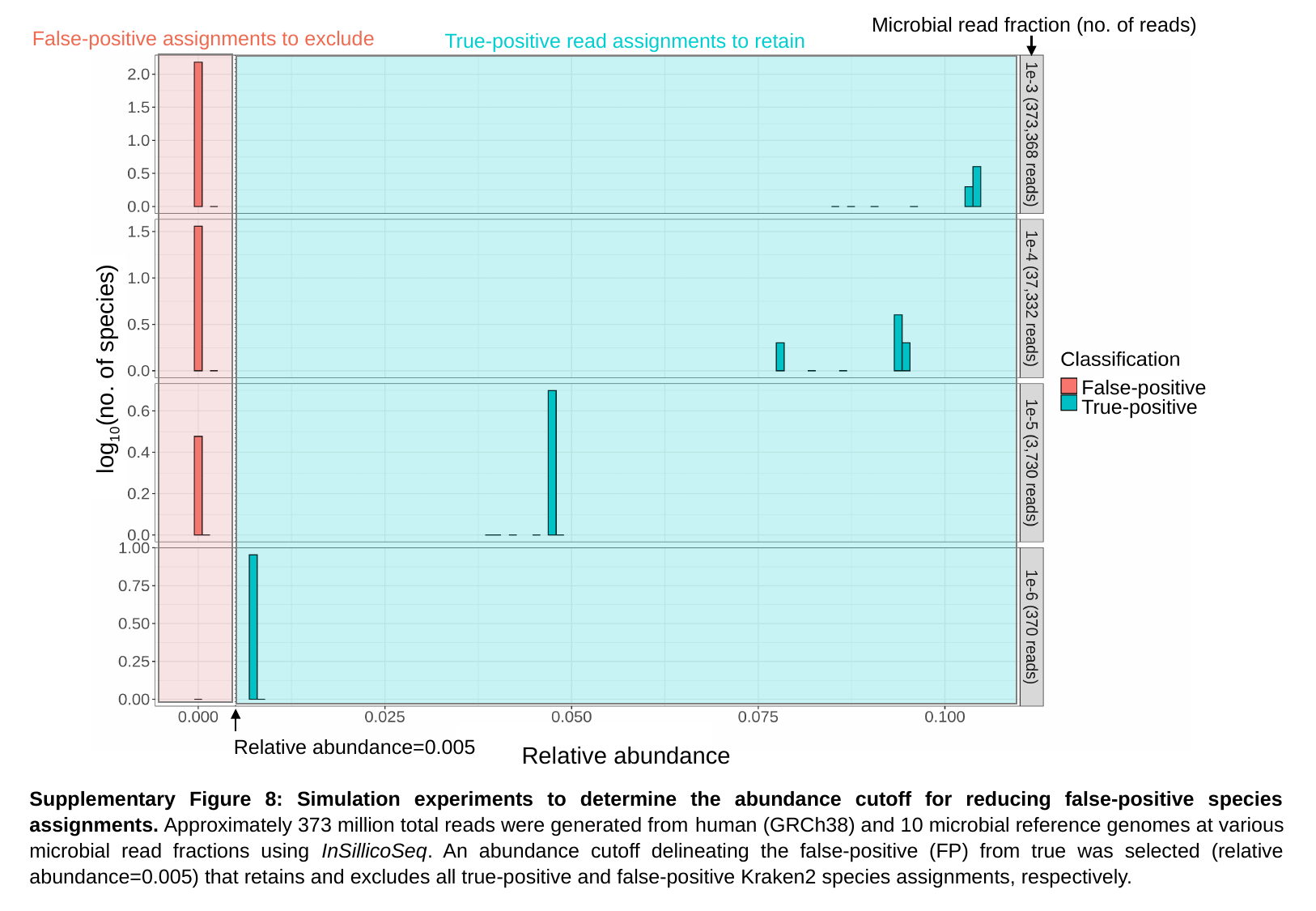

Microbial read fraction (no. of reads)
log_10_(no. of species)
False-positive
True-positive
Relative abundance=0.005
Relative abundance
False-positive assignments to exclude
True-positive read assignments to retain
Supplementary Figure 8: Simulation experiments to determine the abundance cutoff for reducing false-positive species assignments. Approximately 373 million total reads were generated from human (GRCh38) and 10 microbial reference genomes at various microbial read fractions using InSillicoSeq. An abundance cutoff delineating the false-positive (FP) from true was selected (relative abundance=0.005) that retains and excludes all true-positive and false-positive Kraken2 species assignments, respectively.

### Slide 9
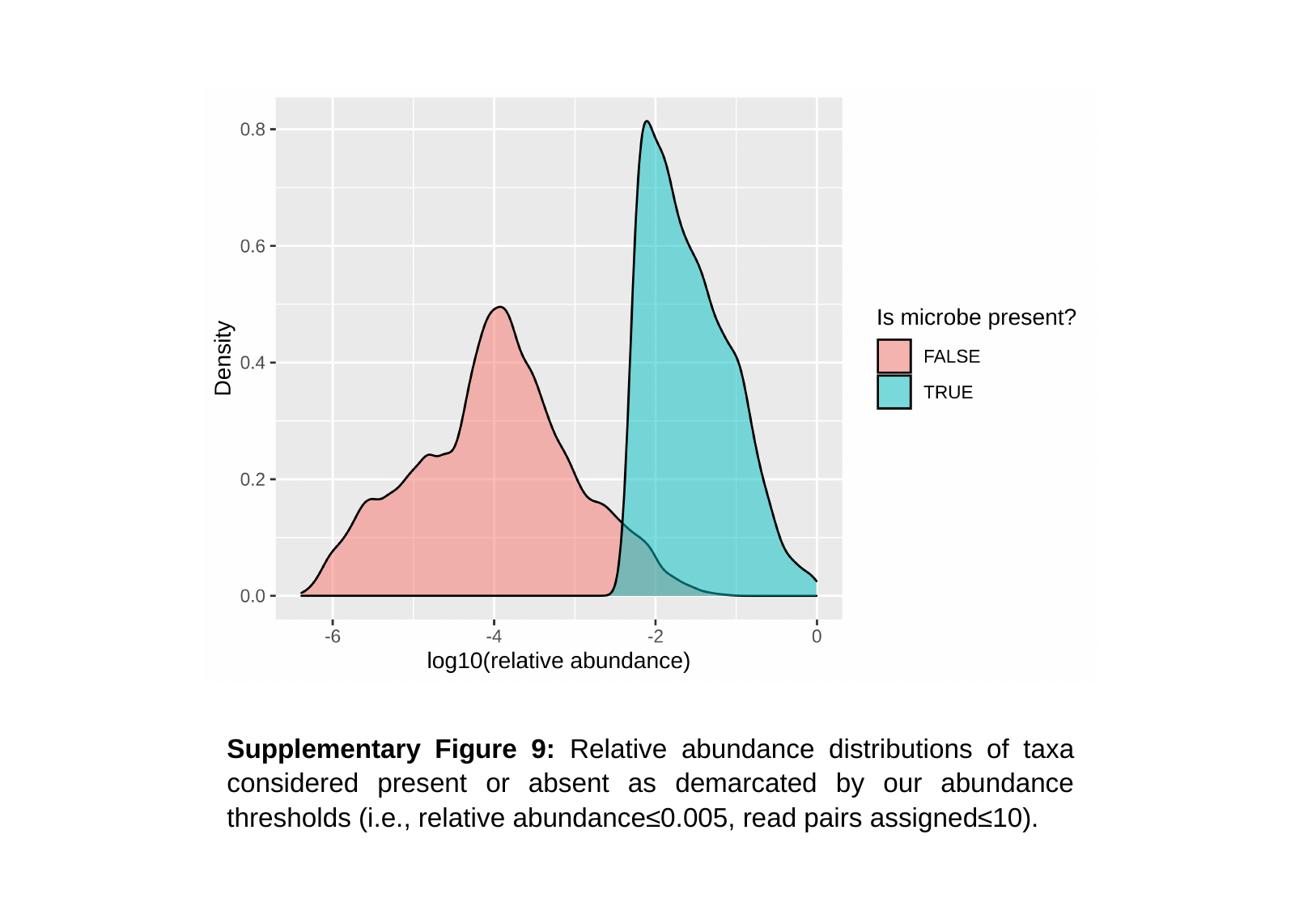

Supplementary Figure 9: Relative abundance distributions of taxa considered present or absent as demarcated by our abundance thresholds (i.e., relative abundance≤0.005, read pairs assigned≤10).

### Slide 10
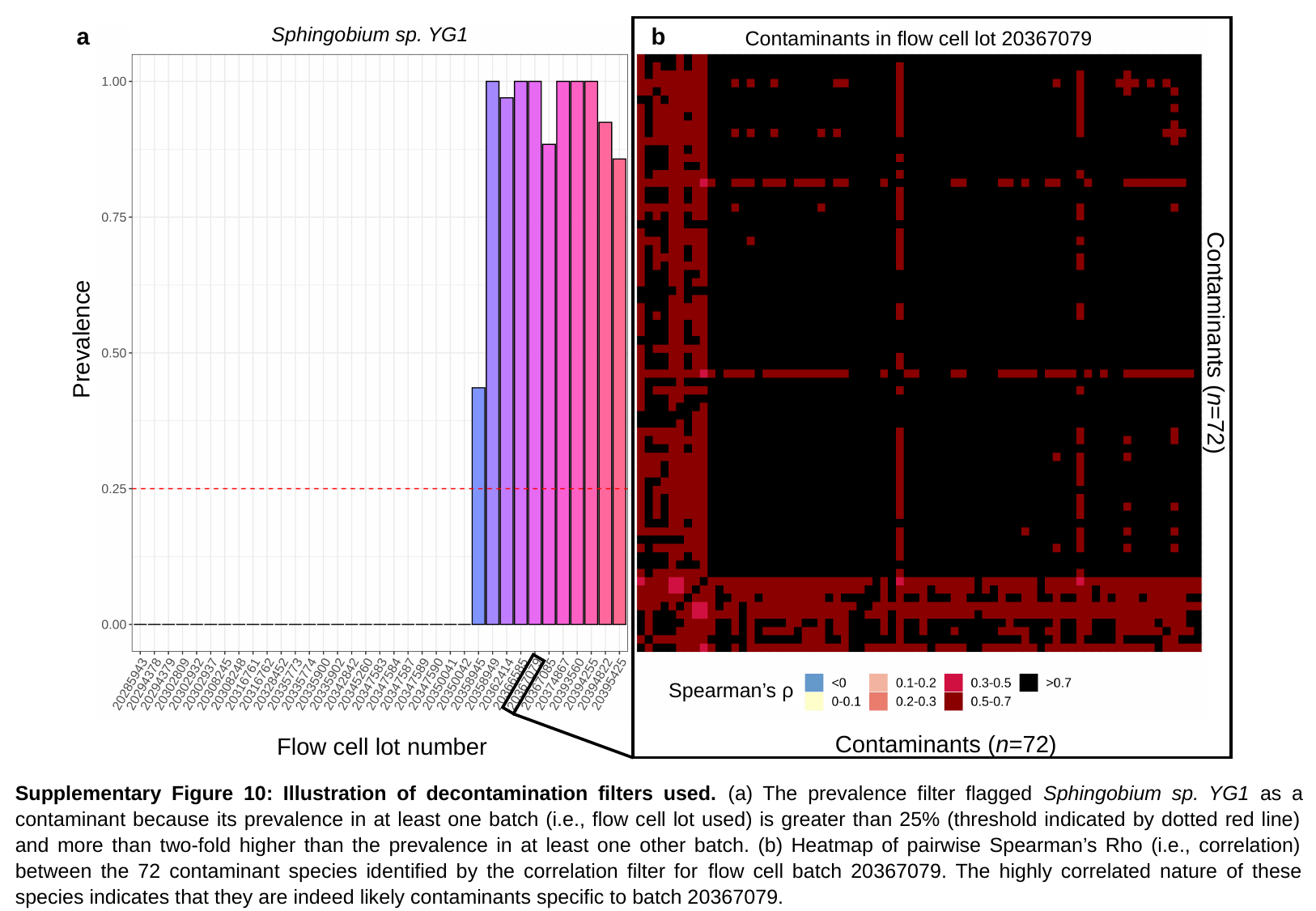

a
Sphingobium sp. YG1
b
Contaminants in flow cell lot 20367079
Prevalence
Contaminants (n=72)
Spearman’s ρ
Contaminants (n=72)
Flow cell lot number
Supplementary Figure 10: Illustration of decontamination filters used. (a) The prevalence filter flagged Sphingobium sp. YG1 as a contaminant because its prevalence in at least one batch (i.e., flow cell lot used) is greater than 25% (threshold indicated by dotted red line) and more than two-fold higher than the prevalence in at least one other batch. (b) Heatmap of pairwise Spearman’s Rho (i.e., correlation) between the 72 contaminant species identified by the correlation filter for flow cell batch 20367079. The highly correlated nature of these species indicates that they are indeed likely contaminants specific to batch 20367079.
